# Quantification of GTPase cycling rates of GTPases and GTPase : effector mixtures using GTPase Glo^™^ assays

**DOI:** 10.1101/2023.11.24.568589

**Authors:** Sophie Tschirpke, Werner K-G. Daalman, Liedewij Laan

**Affiliations:** Bionanoscience Department, Delft University of Technology, Van der Maasweg 9, 2629 HZ Delft, The Netherlands

**Keywords:** Cdc42, GTPase activity, GTPase assay, GTPase effectors, GTPase cycling rates

## Abstract

In different cellular activities like signal transduction, cell division, and intracellular transportation, small GTPases take on a vital role. Their functioning involves hydrolysing guanosine triphosphate (GTP) to guanosine diphosphate (GDP). In this article we explain the application of a commercially accessible GTPase assay, known as the GTPase Glo™ assay by Promega, for the quantitative investigation of GTPase - effector interactions and the interplay between effectors.

**Basic Protocol:** Conducting GTPase assays with GTPase : effector protein mixtures using the GTPase Glo™ assay (Promega).

**Supporting Protocol 1:** Analysing GTPase assays to correlate the assay readout (luminescence) to amount of remaining GTP.

**Supporting Protocol 2:** Fitting GTPase assay data to obtain GTPase cycling rates.

## Introduction

Small Guanosine Triphosphatases (GTPases), are a class of enzymes that play a fundamental role in various cellular processes, including signal transduction, cell division, and intracellular transport. GTPases regulate the hydrolysis of guanosine triphosphate (GTP) to guanosine diphosphate (GDP). Mechanistically, this GTPase activity involves three steps (Fig. 1):(1)binding of GTP to the GTPase, (2) hydrolysis of GTP to GDP and free phosphate, and (3) release of GDP from the GTPase. GTPase activity is often regulated by effector proteins; GTPase activating proteins (GAPs) boost step 2 and GDP/GTP exchange factors (GEFs) enhance step 3 [Cherfils and Zeghouf, 2013, Bos et al., 2009, Vetter and Wittinghofer, 2001]. Several assays exist that examine single GTPase cycle steps, which help us understand the mechanistic details of these cycle steps and their regulation. At the same time, they do not allow studying the interplay of effectors acting on different GTPase cycle steps. Furthermore, approaches that reconstitute the complex cellular functions of GTPases *in vitro* [Vendel et al., 2019, Loose et al., 2008, Bezeljak et al., 2020, Kohyama et al., 2022] are sensitive to variations in protein batch activities and benefit from easy and accessible assays assessing protein purification batches activities.

**Figure 1.**
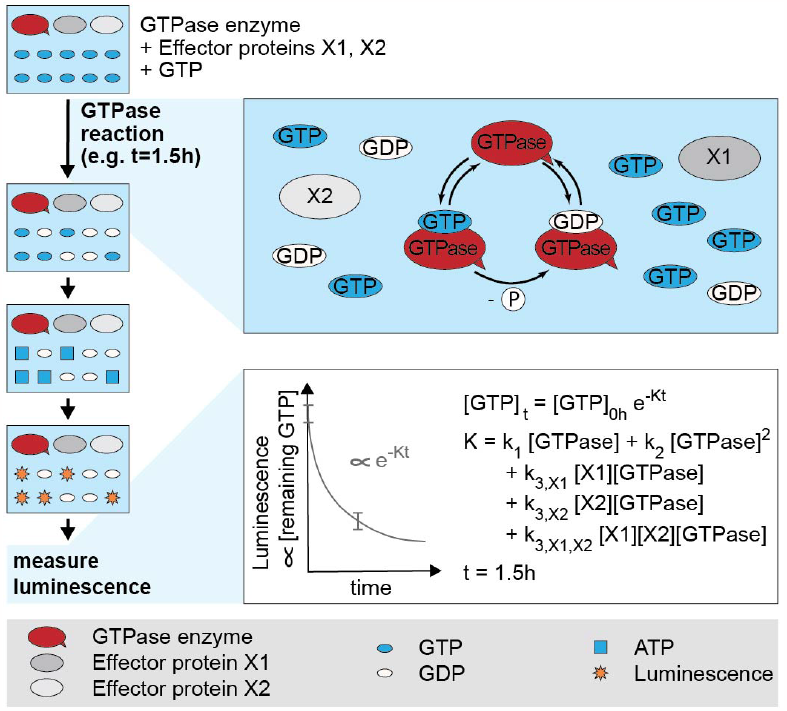
Schematic illustration of the assay steps and GTPase cycling reaction in the GTPase Glo™ assay (Promega). A GTPase, alone or in combination with effector proteins, is incubated with GTP at for a certain time (e.g 1.5 h) in which GTPase cycling occurs. After two processing steps, the amount of remaining GTP, measured as luminescence, is assessed. GTP hydrolysis cycling rates can be extracted by fitting the data with an exponential model. **Abbreviations:** AA amino acid GAP GTPase activating protein GEF GDP/GTP exchange factor

We here describe how a commercially available GTPase assay (GTPase Glo™ assay (Promega), from here on referred to as ‘GTPase assay’) can be used to quantitatively study GTPase effector interactions and effector interplay and to easily activity test GTPase and effector purification batches. In the GTPase assay the proteins of interest are incubated with GTP for a certain amount of time for GTPase cycling to occur. Then the reaction is stopped and the amount of remaining GTP, measured as luminescence signal, is determined. GTPase cycling rates can be obtained from this data through using a coarse-grained exponential fitting model (Fig. 1). This article describes the workflow for conducting GTPase assays using GTPase : effector protein mixtures, and provides analysis software to obtain GTPase cycling rates. We like to note that we do not advice to use this protocol if single GTPase steps are to be examined in-depth, but if effects of a protein on the entire GTPase cycle are of interest. This method is exemplified for a basic GTpase assay using the human Ras (GTPase) (example 1) and for an elaborate GTpase assay set using *S. cerevisae* Cdc42 (GTPase), Cdc24 (GEF), and Rga2 (GAP) [Tschirpke et al., 2023a] (example 2). Tab. 1 summarises how these examples are used to illustrated the protocols described herein. It can also be applied to other GTPases and their effectors. In the Basic Protocol we outline the workflow for conducting GTPase assays using the GTPase Glo™ assay (Promega). In Supporting Protocol 1 we describe how the GTPase assays data (luminescence) can be converted to amounts of remaining GTP. In Supporting Protocol 2 we describe how data obtained by Supporting Protocol 1 can be fitted with a GTPase cycling model (and the by us developed, openly available Matlab code) to obtain GTPase cycling rates.

**Table 1.**
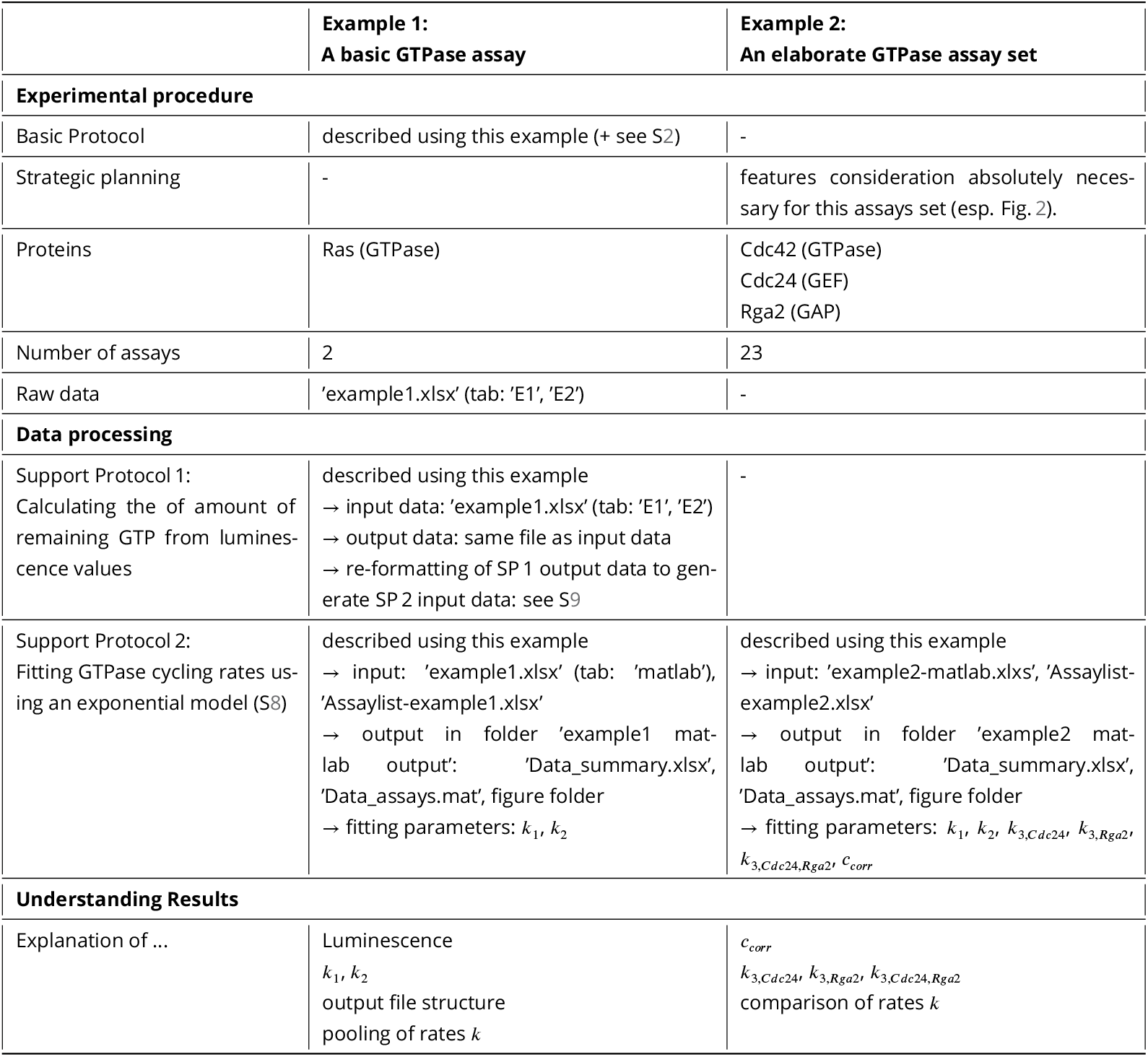
Summary of how the examples are used throughout the protocols. Data files are available at data.4tu.nl. More information can be found in the section: Data Sharing and Data Availability.

|  | **Example 1:**<br>**A basic GTPase assay** | **Example 2:**<br>**An elaborate GTPase assay set** |
| --- | --- | --- |
| **Experimental procedure** |  |  |
| Basic Protocol | described using this example (+ see S2) | - |
| Strategic planning | - | features consideration absolutely necessary for this assays set (esp. Fig. 2). |
| Proteins | Ras (GTPase) | Cdc42 (GTPase)<br>Cdc24 (GEF)<br>Rga2 (GAP) |
| Number of assays | 2 | 23 |
| Raw data | 'example1.xlsx' (tab: 'E1', 'E2') | - |
| **Data processing** |  |  |
| Support Protocol 1:<br>Calculating the of amount of remaining GTP from luminescence values | described using this example<br>→ input data: 'example1.xlsx' (tab: 'E1', 'E2')<br>→ output data: same file as input data<br>→ re-formatting of SP 1 output data to generate SP 2 input data: see S9 | - |
| Support Protocol 2:<br>Fitting GTPase cycling rates using an exponential model (S8) | described using this example<br>→ input: 'example1.xlsx' (tab: 'matlab'), 'Assaylist-example1.xlsx'<br>→ output in folder 'example1 matlab output': 'Data_summary.xlsx', 'Data_assays.mat', figure folder<br>→ fitting parameters: $k_1, k_2$ | described using this example<br>→ input: 'example2-matlab.xlsx', 'Assaylist-example2.xlsx'<br>→ output in folder 'example2 matlab output': 'Data_summary.xlsx', 'Data_assays.mat', figure folder<br>→ fitting parameters: $k_1, k_2, k_{3,Cdc24}, k_{3,Rga2}, k_{3,Cdc24,Rga2}, c_{corr}$ |
| **Understanding Results** |  |  |
| Explanation of ... | Luminescence<br>$k_1, k_2$<br>output file structure<br>pooling of rates $k$ | $c_{corr}$<br>$k_{3,Cdc24}, k_{3,Rga2}, k_{3,Cdc24,Rga2}$<br>comparison of rates $k$ |

### Strategic Planning

If the GTPase assay is done for the first time, we advice to first practice pipetting of the required small volumes into the wells using coloured water. We advice to conduct smaller assays using a larger number of replicas per sample (e.g. 8 samples rows per assay with 5 replicas each, as described in the basic protocol) first, before moving on to larger assay sets with more proteins.

Once you feel more routined with the assay, larger assay sets can be conducted. For example, assays in which the interaction between a GTPase and one or several effectors is studied. The following section applies predominantly (but not solely!) to such larger assays sets involving multiple GTPase enzymes and/or effector proteins (example 2). We advice to determine before starting which assays with how many proteins will need to be carried out. Specifically, we advice to:

#### Verify that the proteins and protein buffers do not interfere with the assay signal

The GTPase assay measures luminescence, which correlates with the amount of remaining GTP. Proteins that interact with luminescence (e.g. fluorescent tags, see S3) and can alter it are therefore not suitable for this assay.

Buffers containing ADP, ATP, GDP, GTP, and guanosine phosphate analogues are also not suitable for the GPase assay, as they interfere with other GTPase assay reactions.

We advice to consider the buffer components and verify that used effector proteins (proteins that are not GTPase enzymes) show no seemingly GTPase activity.

### Verify that the used incubation times lay in the exponential decline regime of remaining GTP

In the GTPase assay the proteins of interest are incubated with GTP for a certain amount of time, in which GTPase cycling can occur. Then the reaction is stopped and the amount of remaining GTP, measured as luminescence signal, is determined. GTPase cycling rates can be obtained from this data through using a coarse-grained exponential fitting model (Fig. 1, S8). The model is based on our observation that the amount of remaining GTP declines exponentially with time (S4). It is still advisable to verified that this is also true for the proteins (GTPase enzymes and GTPase:effector mixtures) and incubation times intended to be used.

To obtain such data, prepare one batch of serial dilutions of the to-be-used proteins. Use exactly these dilutions to conduct several assays using different incubation times. Check if the amount of remaining GTP of these assays follows an exponential decline. Further, analyse each assay individually using our GTPase cycling model. Verify that different incubation times (= different assays) yield the same rates (e.g. S4).

### Dialyse all to-be-used proteins into the same buffer

All proteins should ideally be in the same buffer. In the assay the protein activity is determined by normalising the assay readout (luminescence) of that protein well to that of ‘buffer only’. Differences between those two (in ion concentrations, detergents, or other additives (such as glycerol)) can lead to differences in luminescence. This is easiest avoided if all proteins used in one assay set are in the same buffer. We advice against using a buffer containing glycerol, as the increased viscosity may affect protein activities (unless protein activities in a denser environment are of interest). Further, buffers should *not* contain DTT, GDP, GTP, and other guoanosine analogues, ADP, ATP, and other nucleotide triphosphates, EDTA or other chelating agents that can complex magnesium. If a GTPase GEF interaction ought to be investigated, the buffer usually requires magnesium salts (e.g. 10 mM MgCl_2_).

Many proteins require glycerol in the buffer for storage. We advice to keep the protein at a high concentration in a buffer containing glycerol (e.g. 50 mM Tris-HCl (pH=7.5), 100 mM NaCl, 10 mM MgCl_2_, 1 mM 2-mercaptoethanol, 10% glycerol), and then dilute it into a buffer of the same composition *without* glycerol (e.g. 50 mM Tris-HCl (pH=7.5), 100 mM NaCl, 10 mM MgCl_2_, 1 mM 2-mercaptoethanol). GTPase assays usually require small concentrations of proteins (in our experience 0.5-5 µM). If the protein is stored at a high concentration, the amount of glycerol in the final sample in the GTPase reaction will be negligible.

If it’s not possible to transfer all proteins into the same buffer, it needs to be verified that the differences in buffer composition do not affect luminescence intensities. A GTPase assay containing only the different buffer mixtures can elucidate this: If all buffer mixtures exhibit the same luminescence values then the difference in buffer composition do not affect luminescence. However, it is still possible that the buffer components affect protein behaviour!

#### Use the same serial dilution for all assays

The assay is sensitive to concentration changes (especially of highly active GPases and of effectors strongly boosting GTPase activity), and thus sensitive to any pipetting errors when preparing the protein dilutions. We advice to prepare serial dilutions of every protein, and use exactly the same serial dilution for the entire assay series. Different serial dilutions of the same protein can exhibit slightly different rates. For example, if the effect of effector protein X on GTPase A and GTPase B ought to be compared, the same serial dilution of effector X ought to be used for both GTPases.

#### Include a reference sample if data from different assays should be compared

GTPase assays can be sensitive to GTPase activity changes, which can be due to small pipetting errors or condition changes (incl. temperature, shaker speed, …). This can lead to slight changes in protein activity between separate assays. It is therefore best to include all samples that ought to be compared into the same assay.

However, this is not always possible. If the strength of several effectors on one GTPase enzyme ought to be compared, we advice to always include one sample containing only the GTPase into each assay (see Fig. 2). Then the assay-specific activity of the GTPase sample can be used to account for any assay-specific variability (through fitting the correction factor *c*_*corr*_, as will be explained in later sections and in S8 Eq. 9).

**Figure 2.**
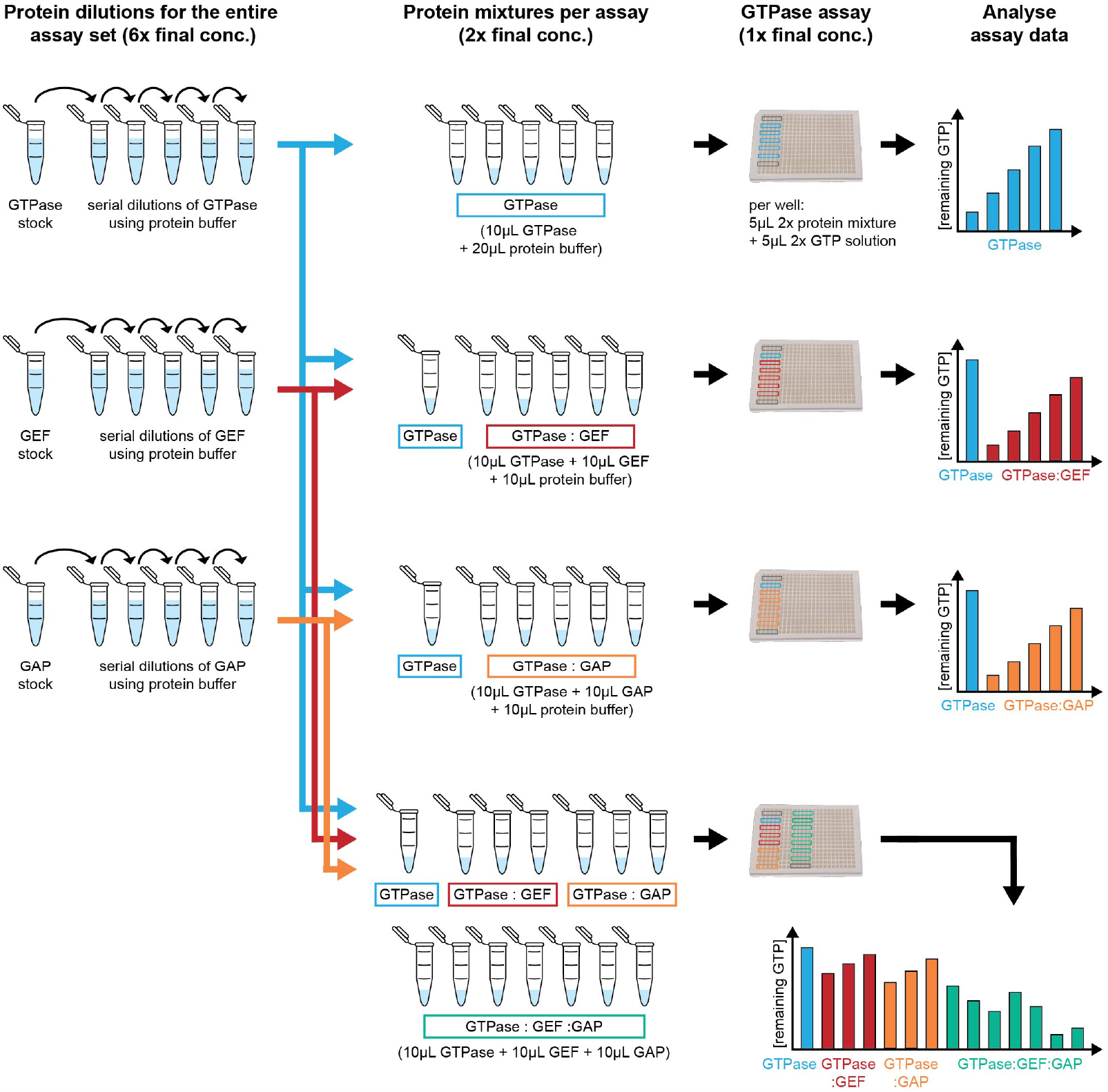
Schematic illustrating the assays and protein dilutions required for conducting a GTPase assay set investigating GTPase, GEF, and GAP actitvity and GEF-GAP interaction (example 2).

(Usually these assay-variations are very small (*c*_*corr*_ ≈ 1.0) [Tschirpke et al., 2023b, Tschirpke et al., 2023a]. We advice to exclude assays with a big or very small *c*_*corr*_, as these indicate that the GTPase behaviour/assay conditions are unusual.)

### Determine useful concentration ranges before conducting an assay series

In our experience a good readout regime is between 5-90% remaining GTP. Higher values (90-100% remaining GTP) have a bigger error (due to the normalisation), and in lower regimes (0-5% remaining GTP) saturation effect due to little remaining GTP might take place. In this regime a saturation of the fitted cycling rates may occur. If an assay with multiple proteins (e.g. a GTPase and two effector proteins) is conducted, it is useful to determine beforehand which concentration ranges of each protein lead to optimal readouts to reduce the time it takes to conduct the entire assay series.

It is also advisable to stick to similar incubation times for all assays, as this streamlines the selection of protein concentrations (for assays containing protein mixtures) that yield suitable readout regimes.

### Make an assay plan

If multiple protein interactions ought to be quantified and compared, the same serial dilutions of these proteins ought to be used. Given that we advice against the use of a glycerol-containing buffer, that many proteins require glycerol when being frozen, and that many proteins are not stable for weeks at room temperature or 4°C, all required assays need to be conducted within a rather short time period to obtain reliable and comparable results. We therefore strongly recommend to carefully plan which assays with which proteins need to be conducted, and calculate the required volumes of protein needed.

We will explain our recommended procedure on the example of a GTPase assay series that investigates the GTPase cycling of a GTPase, and the effect of both a GEF and a GAP, alone and in combination, on it (example 2, Fig. 2) [Tschirpke et al., 2023a]:

#### Considerations

(as we described earlier): (1) Different serial dilutions of the same protein might exhibit slightly different rates (due to small pipetting errors when making the dilution). Hence, the same dilution should be used for the entire assay set. (2) For larger assays it is advisable to already know which concentration ranges give good signal to reduce the time the proteins need to be stored. (3) Include a reference sample (here: sample of the GTPase enzyme alone in the concentration used in the particular assay). (4) The incubation times used are in the exponential regime. (5) It is advisable to use similar incubation times for all assays, as this will make it easier to choose protein concentrations for assays containing protein mixtures that will result in suitable readout regimes.

#### Protein dilutions

To conduct the assay, several protein solutions need to be made: Protein dilutions of GTPase, GEF, and GAP that will be used to conduct the entire assay set (Fig. 2 left column). These starting dilutions will be used to create protein mixtures, that will be used to for one GTPase assay (Fig. 2 middle column).

Which concentrations should these starting dilutions have? To initiate the GTPase reaction, a 2× GTP solution is mixed in a 1:1 volume ratio with the protein mixtures. Therefore, the protein mixture needs to be 2× of the final concentration in the assay. Further, in the final assay the protein mixture needs to contain the GTPase, GEF, and GAP. It is easiest to make this mixture if all proteins can be mixed at 1:1:1 volume ratios. Hence, the initial starting protein serial dilutions, which will be used for all assays of the set, should be 6× stocks of the final protein concentrations. (Similarly, if an assay set will contain at maximum a mixture of 4 (5) proteins, the initial starting protein serial dilutions need to be 8× (10×) stocks of the final protein concentrations.

#### Procedure

In which order should the assays be conducted? First, a GTPase assay with a serial dilution of only the GTPase enzyme is conducted (Fig. 2 top row). Pick one or two GTPase concentrations to conduct all following assays with. In this case the following assays will involve a GEF and a GAP, both of which boost the activity of the GTPase. It is therefore wise to choose a GTPase concentration that leads to a high amount of remaining GTP (e.g. 90-80% remaining GTP). Next, a GTPase assay with the GTPase and several GEF concentrations (obtained from serial dilutions) is performed. Use the same GTPase concentration for all samples; samples of GTPase:GEF mixtures, and a sample containing only the GTPase (Fig. 2 second row). The same procedure is applied to the GAP: Conduct a GTPase assay with the GTPase and several GAP concentrations (obtained from serial dilutions). Use the same GTPase concentration for all samples; samples of GTPase:GAP mixtures, and a sample containing only the GTPase (Fig. 2 third row). Lastly, perform GTPase assays in which all proteins are present (GTPase alone, GTPase:GEF mixtures, GTPase:GAP mixtures, and GTPase:GEF:GAP mixtures, all using the same GTPase concentration). Choose GEF and GAP concentrations that resulted in a medium amount of remaining GTP (e.g. 60-70% remaining GTP), to leave room for even lower values when both proteins are combined (Fig. 2 bottom row). Ideally additional assays will be added to control for potential non-canonical effects (see below).

##### Control for non-canonical effects through adding an inert protein

We observed that the non-canonical effects can take place during GTPase cycling in the GTPase assay. These effects are generally small, and can be accounted for: For example, we observed that both Bovine serum albumin (BSA) and Casein slightly boost GTPase activity of both Ras and Cdc42. BSA and Casein are considered inert and have no known interaction with neither Ras or Cdc42. We suspect that they seemingly boost the GTPase’s activity through increasing the effective GTPase concentration: A few GTPase molecules might stick to the well wall in each reaction chamber, being rendered inactive. Through adding another protein the chamber wall will be covered with both some GTPase molecules and some molecules of the other protein, thereby increasing the effective GTPase concentration.

To ensure that the effect of a protein on GTPase cycling is not only due to these non-canonical effects, we advice to additionally conduct assays with the GTPase and an inert protein (e.g. BSA or Casein). The effect of the to-be-studied protein will need to exceed that of a equimolar concentration of the inert protein to be of non-canonical origin.

Both BSA and Casein are considered inert proteins. However, BSA is strongly negatively charged, which could lead to non-specific protein interactions. Therefore Casein, or another supposedly inert protein, might be more suitable as a control. For the previously used example 2 (an assay series including a GTPase, GEF, and GAP, Fig. 2), control assays for non-canonical effects include: (1) GTPase:Casein serial dilution, (2) GTPase:GEF:Casein, (3) GTPase:GAP:Casein.

### Basic Protocol: Conducing GTPase assays

In the GTPase assay the proteins of interest (GTPase enzymes with or without effectors) are incubated with GTP for a certain amount of time for GTPase cycling to occur. Then the reaction is stopped. The remaining GTP is, through two follow up reactions, translated into luminescence signal, which is measured (Fig. 1).

The following protocol steps are, with only minor modifications, those described in the assay manual. The volumes are for assays conducted in 384-well microplates. Assays using larger volumes can be conduced using different plates (see GTPase Glo™ assay manual). In addition to following the steps described here we advice to carefully read the assay manual [Promega Corporation, 2015].

The following protocol describes the steps to conduct a GTPase assay using 6 concentrations of Ras GTPase (example 1) (Fig. 3). An illustration of required assay planning steps is shown in S2. The assay results in data found in ‘example1.xlsx’ (tab: ‘E1’, ‘E2’).

**Figure 3.**
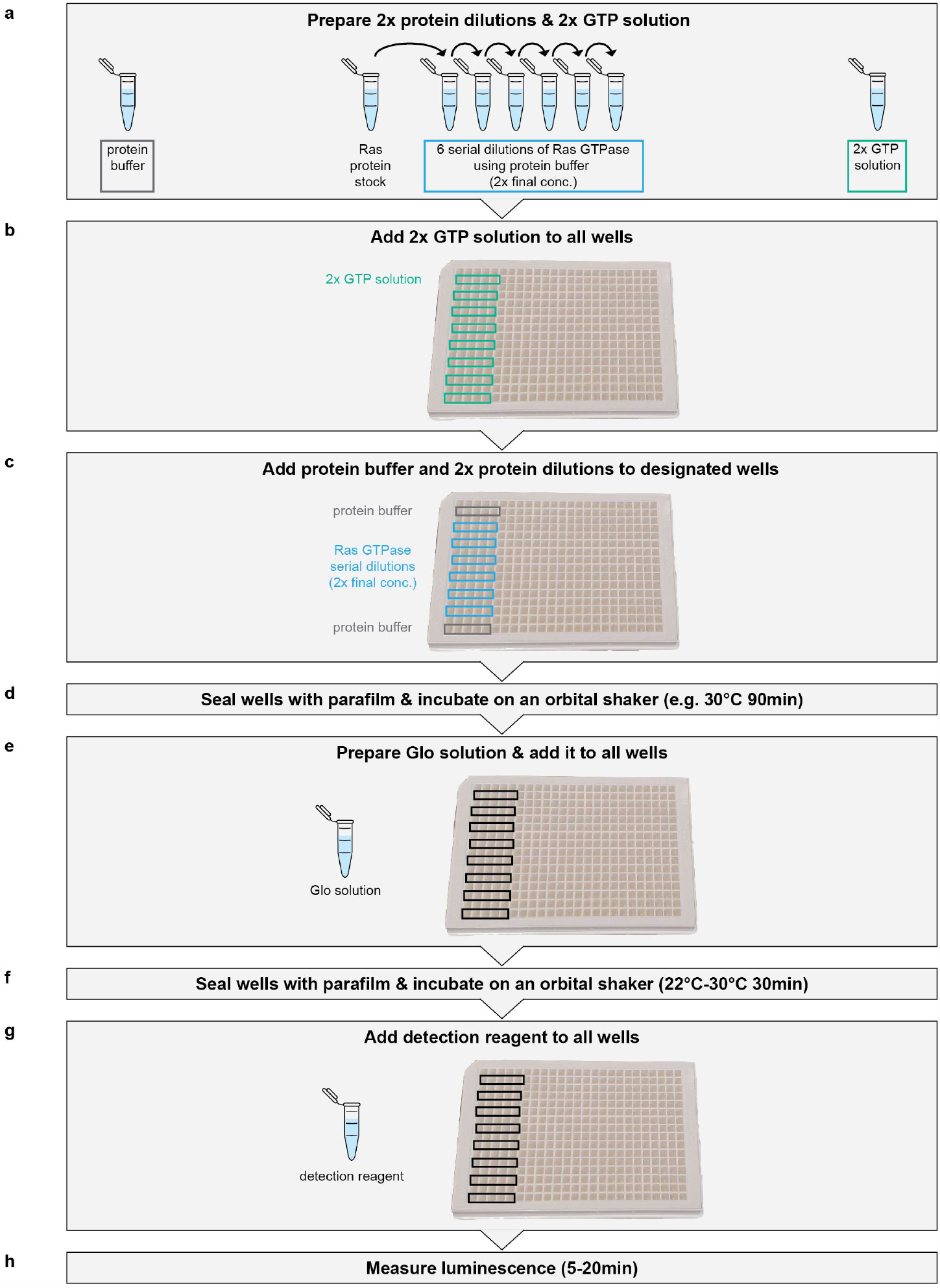
Schematic illustration the steps necessary to conduct a GTPase assay using a serial diluton of 6 Ras GTPase concentrations. The preparatory steps to conduct this assay are illustrated in S2.

#### Materials

- GTPase Glo™ assay (Promega, cat. no. V7681 (1’000 reactions) or V7682 (10’000 reactions))
- proteins of interest: a GTPase enzyme (and optionally effector proteins). As a test GTPase human Ras (EMD Millipore, cat. no. 553325) can be used.
- buffer in which the proteins of interest are stored (’protein buffer’)
- optional: Casein (Sigma-Aldrich, cat. no. C7078) or Bovine serum albumin (BSA) (Thermo Scientific, cat. no. 23209)
- 1.5 mL and 15 mL reaction tubes 384-well white flat bottom microplates (Corning, cat. no. 3572)
- parafilm (Bemis Company Inc., cat. no. PM996)

### Equipment

- pipettes: 0.5 µL, 10 µL, 20 µL, 100 or 200 µL, 1 mL
- vortex mixer (e.g. Thermo Scientific, cat. no. 88882011)
- Mini centrifuge (e.g. Thermo Scientific, cat. no.75004061)
- orbital shaker (e.g. Innova 2300 platform shaker (New Brunswick Scientific))
- optional: temperature-controlled incubator
- a plate reader that can measure luminescence (e.g. Synergy HTX plate reader from BioTek)

#### Preparation of proteins

1. Dialyse the GTPase enzyme into a suitable buffer (’protein buffer’). *E*.*g. 50* mM *Tris-HCl (pH=7*.*5), 100* mM *NaCl, 10* mM *MgCl*_2_, *1* mM *2-mercaptoethanol. More considerations on the buffer are described in the strategic planning section*. *If the GTPase enzyme is at a high concentration (e*.*g. 50-100* mM*) and the buffer the protein is stored in does not differ that much from the protein buffer, it can be sufficient to only dilute the protein using protein buffer*.

#### Before starting the assay

2. ’Mise-en-place’: Decide on the assay scope and prepare all assay materials needed. Templates can be found in S1. An example for an assay of Ras GTPase is shown in S2 and Fig. 3.
  a. Decide which samples and how many replica wells per sample you want to use. *We recommend 3-5 replicas per sample. In the beginning In the beginning we recommend using one GTPase enzyme per assay and 5 replicas. We suggest using 8 samples per assay: 6 concentrations of the GTPase enzyme (here: Ras GTPase) + 2 protein buffer samples (for normalisation) (Fig. 3)*.
  b. Calculate which volumes of protein dilutions, 2× GTP solution, Glo solution, and detection reagent are needed to conduct one assay (example calculation are given in S2). Vol.(protein dilution) = 5µL × number of replicas Vol.(2× GTP solution) = 5µL × number of replicas per sample × number of samples Vol.(Glo solution) = 2 × Vol.(2× GTP solution) Vol.(detection reagent) = 4 × Vol.(2× GTP solution) *We recommend to always prepare a bit more of the solutions that needed*. *We recommend preparing 5*µL *more of each protein dilution than calculated, to ensure that there is also sufficient solution for the last well. E*.*g. for 5 replicas per sample, for each dilution prepare 5*µL × *5 + 5*µL *= 30*µL.
  c. Calculate the dilution series steps for proteins used. Note that the solutions prepared need to be 2× of the final concentration in the assay (example calculation are given in S2).
  d. Prepare the plate: seal the assay wells that will not be used with parafilm to prevent contamination with dirt, and label/mark the wells where sample will be added. *Leave one empty row between all sample rows to avoid any spill-over of luminescence signal between samples (see Fig. 3 and S5)*.
  e. Cut 4 pieces of parafilm that are sufficiently big to cover the area where samples will be added.

#### Conducting GTPase Glo™ assays

3. Thaw protein samples and assay solutions on ice.
4. Serial dilute the Ras GTPase enzyme with protein buffer (according to your calculation) to obtain 2× GTPase enzyme dilutions (S2 and Fig. 3a). *Vortex for proper mixing, then spin down mildly to collect all volume in the bottom of the tube*.
5. Prepare a 2× GTP solution. A pipetting scheme is given in Tab. 2. *The 2*× *GTP solution should always be freshly prepared immediately before starting the assay. Vortex the GTP and DTT before use. Vortex the 2*× *GTP solution for proper mixing. The 2*× *GTP solution can only be used freshly and should not be stored/frozen (S6). We advice to aliquot GTP and DTT to reduce the number of freeze-thaw cycles, in our experience up to 3 freeze-thaw cycles do not impact assay performance*.
6. Add 5µL 2× GTP solution to all wells (Fig. 3b).
7. Add 5µL protein buffer and 2× Ras GTPase dilutions to designated wells (Fig. 3c). *We advice to have one row containing protein buffer on top of the assay and one row in the bottom. In rare cases the luminescence values slightly drift towards lower values throughout the assay. Placement of a buffer row as a first and last sample row ensures that this drift can be (1) detected and (2) accounted for in the analysis*.
8. Seal the wells with two sheets of parafilm and incubate for a designated time on an orbital shaker. *We recommend to seal the wells through placing one sheet of parafilm on top of the plate (without stretching it) and firmly pressing it into to plate with a rounded surface (such as sissor handles). Then repeat the same process with the second sheet of parafilm. This ensures that the wells are tightly closed. Using only one sheet of parafilm can result insufficient sealing (small holes in the parafilm from the too much pressure)*. *The incubation time and temperature depends on the protein of interest and used concentrations. We recommend choosing a temperature in accordance to the protein’s environment in vivo. We recommend incubation times of 60-90 min*.
9. Prepare the Glo solution a few minutes before the incubation time is reached. A pipetting scheme is given in Tab. 3.Then add 10µL Glo solution to all wells (Fig. 3e). *Preparation of the Glo solution:*
  a. *Vortex the Glo buffer and 10* mM *ADP before use. Do NOT vortex the GTPase-Glo reagent. Instead, gently tap it for proper mixing*.
  b. *To use the pipetting scheme provided in Tab. 3, 10* mM *ADP provided in the assay kit needs to be diluted to 1* mM *ADP using ultra pure water!*
  c. *We advice to aliquot 10* mM *ADP, Glo buffer, and Glo reagent to reduce the number of freeze-thaw cycles. These aliqouts can be re-used (S6)*.
  d. *Both 1* mM *ADP and Glo solution can only be used freshly and should not be stored/frozen. Vortex both for proper mixing*.
  e. *The Glo reagent is the reagent that usually runs out first. To increase the number of assay runs per kit, prepare only the volume needed*.
10. Seal the wells with two sheets of parafilm and incubate for 30 min on an orbital shaker.
11. Add 20µL detection reagent to all wells (Fig. 3g). *We advice to aliquote the detection reagent to reduce the number of freeze-thaw cycles and decrease the time required for thawing*. *Calculate how much detection reagent is required per assay run. In some cases we observed that two seperately stored aliquots from the same batch can lead to distinct luminescence values (S7). To avoid such a shift within an assay, pull a sufficient volume into one tube (e*.*g. pull several aliquots) before adding the detection reagent to the wells. Vortex for proper mixing*.
12. Measure the luminescence of each well using a plate reader for 20 min. *Activate orbital shaking between measurements to ensure proper mixing. If the plate reader does not have a shaking option, incubate the plate for 5 min on an orbital shaker before measuring luminescence. Then measure for 15 min*.

### Support Protocol 1: Basic analysis of GTPase assay data

This protocol describes basic analysis steps to calculate from the GTPase assay readout (luminescence) the amount of remaining GTP. The amount of remaining GTP is a simple measure used to compare the GTPase activity of different GTPases (of the same concentration, incubated with GTP for the same amount of time).

The protocol illustrates the steps for GTPase data from example 1: an assay including 6 Ras GTPase serial dilutions and two buffer reference samples, with each 3 replicas per sample (see basic protocol and S2). This protocol uses *and* generates data in ‘example1.xlsx’, tab: ‘E1’, ‘E2’.

#### Necessary resources

- A spreadsheet editor or other analysis software (e.g. python script)

#### Protocol steps

1. Calculate the average luminescence for time points 5-20 min per well (= replica) (Fig. 4)
2. Calculate the luminescence average for each sample: Average between wells (=replicas) of the same sample (Fig. 5a). *Combine the wells/replicas of the first and last buffer row. Calculate the average using all values. I*.*e. do not distinguish between the first and last buffer row (see Fig. 3, S2)*.
3. Calculate the standard error of the mean for each sample (Fig. 5b).
4. Use the average luminescence values (Fig. 5a) to translate luminescence into amount of remaining GTP using Eq. 1 (Fig. 5c): The luminescence values of the buffer samples correspond to 100% remaining GTP. The amount of remaining GTP of each protein sample corresponds to the luminescence of this sample divided by the luminescence of the buffer. *This step normalises the luminescence values of the protein samples to 100% remaining GTP (luminescence of the buffer). In principle, the protein samples also need to be normalised to 0% GTP. Given that we observed basically no luminescence in the blank samples (S5), this step is not necessary*.

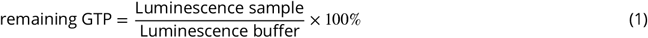
5. Calculate the error (‘Δ remaining GTP’) using error propagation (Eq. 2, Fig. 5d): ‘Lum.’ is the average luminescence and ‘ΔLum.’ refers to the standard error of the mean.

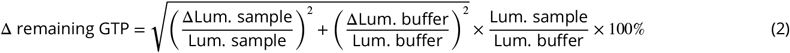

The following step is only required if the data analysis will be continued using Supporting Protocol 2:
6. Re-format the data in a spreadsheet to follow the formatting shown in Fig. 6. This can be done using the python script provided in S9. In short, the data must be organised into the following columns with specific headers:
  - ’Run’: state the assay number. The assay number must start with an ‘E’ followed by a number, which can be followed by a letter (and underscore letter). Eg. E1, E2, E100, E1a, E100abc, E1_a, E100_abc, E1a_abc, E100f_abc
  - ’Time’: incubation time, stated in hours
  - ’GTP_remaining’: amount of remaining GTP, normalised to 1
  - ’Error’: error values, normalised to 1
  - ’Buffer_error’: error of the buffer, normalised to 1
  - ’[GTPase]_conc’: The header must contain the name of the GTPase, followed by ‘_conc’ (e.g. Ras_conc, Fig. 6). The concentration must be stated in µM
  - optional: ‘[Effector]_conc’: The header must contain the name of the effector, followed by ‘_conc’. The concentration must be stated in µM. This column should only be there if it contains more than one unique value per assay. *We advice to group assays of the same type (i*.*e. containing the same proteins and dilution types) into one speadsheet tab. They must follow the same structure. The header can only be included once. E*.*g. An assay set examines the GTPase Cdc42 and its GEF Cdc24. Two types of assays were conducted: (A) GTPase assays using only serial dilutions of the GTPase Cdc42. (B) GTPase assays using a constant Cdc42 concentration and serial dilutions of the GEF Cdc24. We advice to group all assay data of (A) into one tab and all assay data of (B) into one tab*. *More examples are given in Support Protocol 2*.

**Figure 4.**
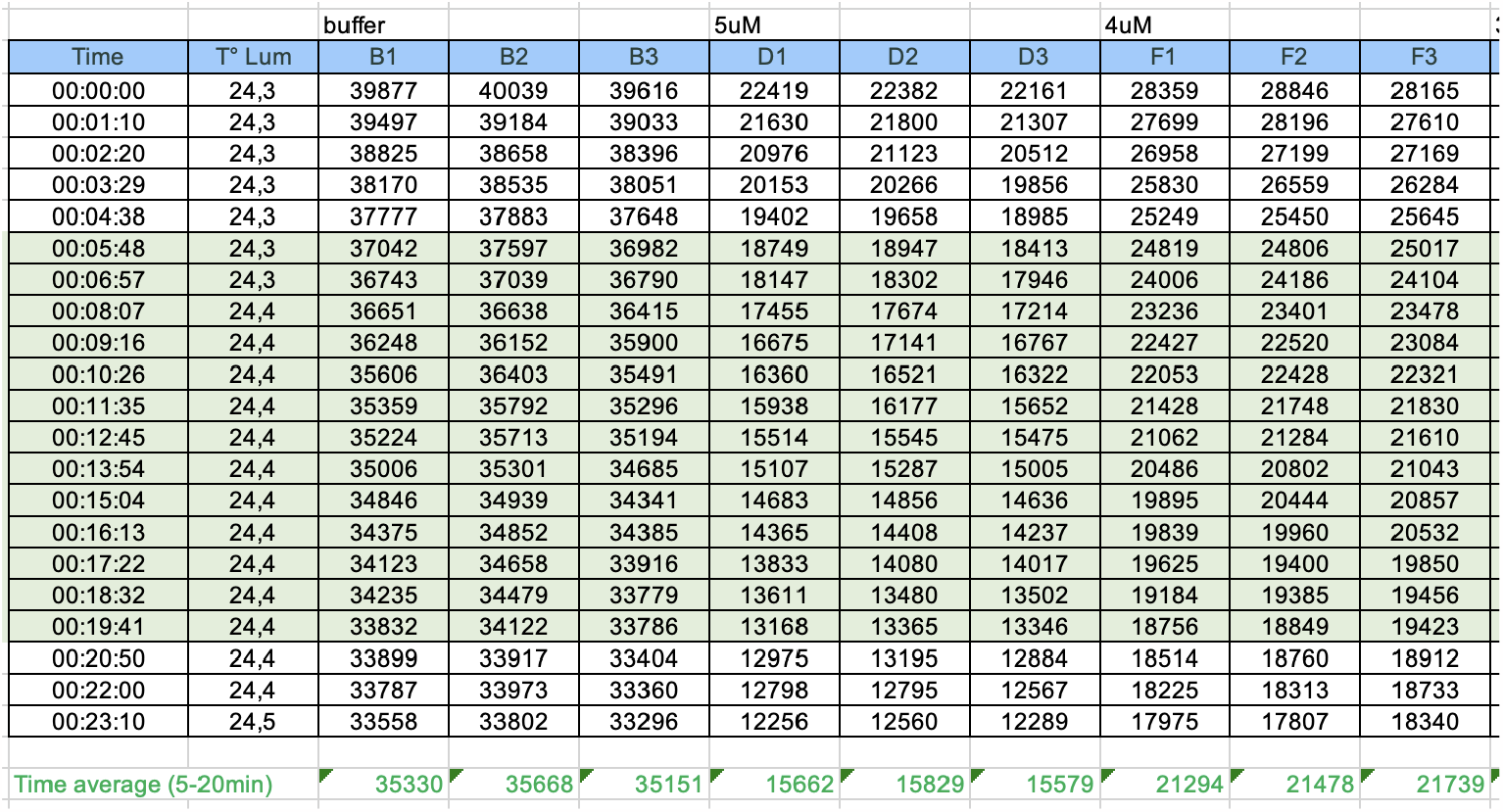
Data analysis part 1 for the GTPase assay described in the basic protocol (Fig. 3 and S2): Time-averaging luminescence values. (Data is taken from ‘example1.xlsx’, tab: ‘E1’.)

**Figure 5.**
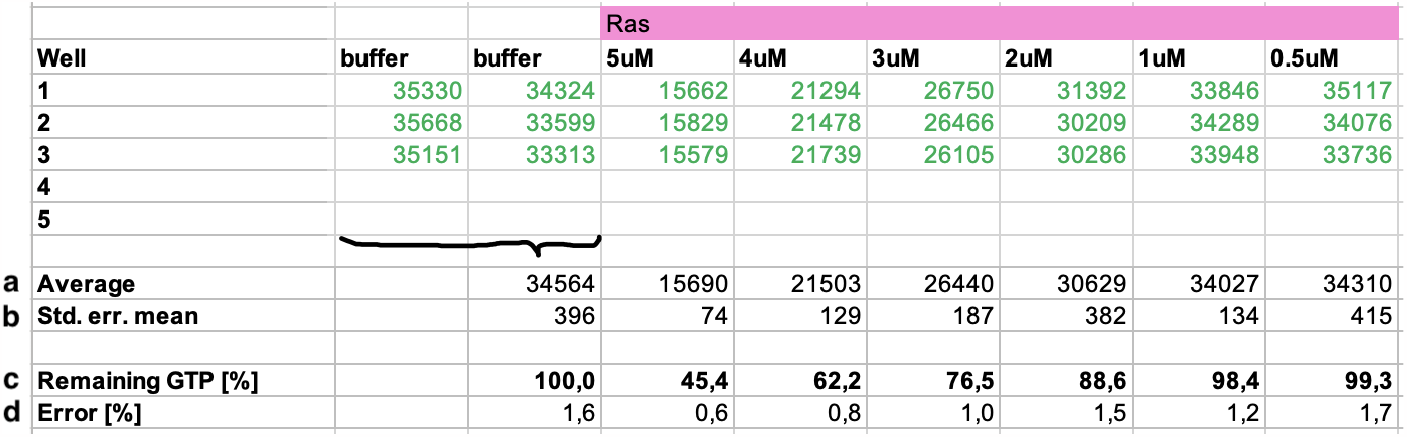
Data analysis part 2 for the GTPase assay described in the basic protocol (Fig. 3 and S2): calculating the amount of remaining GTP from luminescence values. (Data is taken from ‘example1.xlsx’, tab: ‘E1’.)

**Figure 6.**
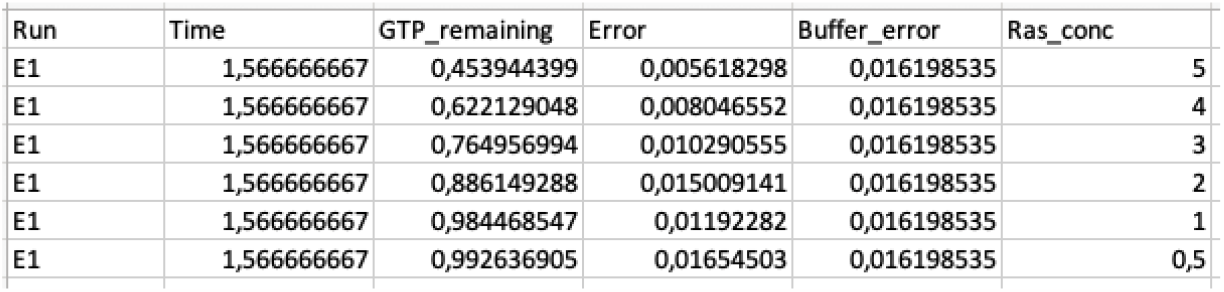
Reformatting of data Fig. 5 (e.g. using a python script provided in S9). (Data is taken from ‘example1.xlsx’, tab: ‘E1’.)

**Table 2.** Pipetting scheme for the 2× GTP solution.

| Starting solution | final concentration | 500 µL | 750 µL | 1 mL |
| --- | --- | --- | --- | --- |
| protein buffer | - | 494 µL | 742 µL | 989 µL |
| DTT (100 mM) | 1 mM | 5 µL | 7.5 µL | 10 µL |
| rGTP (10 mM) | 10 µM | 0.5 µL | 0.75 µL | 1 µL |

**Table 3.**
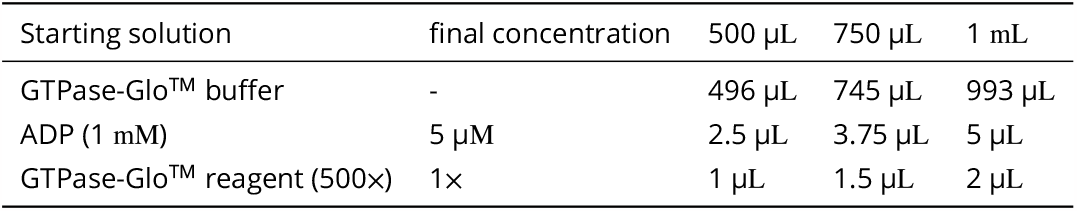
Pipetting scheme for the Glo solution. Please note, to follow this pipetting scheme the ADP solution provided in the kit (ADP, 10 mM) has to be diluted to 1 mM (e.g. mix 2 µL ADP (10 mM) with 18 µL of ultra pure water)!

| Starting solution | final concentration | 500 µL | 750 µL | 1 mL |
| --- | --- | --- | --- | --- |
| GTPase-Glo™ buffer | - | 496 µL | 745 µL | 993 µL |
| ADP (1 mM) | 5 µM | 2.5 µL | 3.75 µL | 5 µL |
| GTPase-Glo™ reagent (500×) | 1× | 1 µL | 1.5 µL | 2 µL |

### Support Protocol 2: Fitting GTPase cycling rates

This protocol describes how the data, pre-processed using the steps described in Support Protocol 1, can be fitted using a GTPase activity model (S8) to obtain GTPase cycling rates. It can be used to fit data of GTPases and GTPase - effector *X* mixtures (up to 2 effectors).

In short, first data of GTPase serial dilutions are fitted with an exponential to obtain GTPase cycling rates *k*_1_ and *k*_2_:

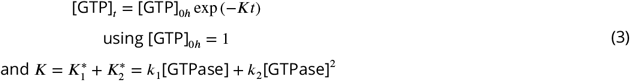

Here *K* refers to the ‘overall GTP hydrolysis rate’ and depends on the concentration of the GTPase. We call the concentration-independent rates *k* GTPase cycling rates, referring to the fact that they describe GTPase cycling (and not specific GTpase cycle steps). The overall hydrolysis rate *K* is composed of an unaided hydrolysis contribution by an individual GTPase 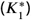, and an cooperative contribution by two GTPase molecules 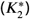.

Then, mixtures of one GTPase and one effector *X*_1_ are fitted with

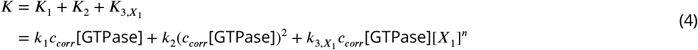

and mixtures of one GTPase and two effectors (*X*_1_, *X*_2_) are fitted with

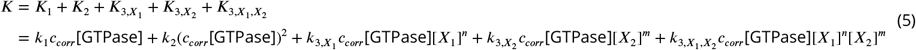

using *n* and *m* either 1 or 2. This fits yield correction factors *c*_*corr*_ and cycling rates 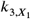 (and 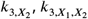 ). *c*_*corr*_ is dimensionless and accounts for variability between assays. It ought to be close to 1.0 (S8). *K*_1_ and *K*_2_ represent corrected version of 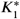 and 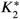 obtained from the GTPase serial dilutions 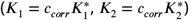. 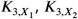, and 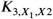 represents the overall hydrolysis rate contribution connected to effectors. They depend linearly on the corrected GTPase concentration, but also on the respective effector concentrations, either linearly or quadratically.

Note, in order to analyse GTPase - effector mixtures, *k*_1_ and *k*_2_ must be known. It is thus required to fit data of GTPase (serial) dilutions first. The model allows to fit GTPase - effecotr mixtures of up to two effectors present, with each effector showing either a linear or quadratic concentration dependence. If the effectors show neither of these concentration-dependencies (e.g. due to saturation), we advice to either only include the linear/quadratic regimes into the analysis or extend our fitting model to match the specific case. With the provided software it thus is possible to fit the following protein (mixtures):

- GTPase serial dilution (example 1)
- GTPase + effector serial dilution (linear or quadratic conc. dependence, Eq. 4) (example 2)
- GTPase + effector *X*1 serial dilutions + effector *X*2 serial dilutons (Eq. 5, Fig. 1) (example 2)

The model is based on our observation that the amount of remaining GTP declines exponentially with time (S4). It is still advisable to verified that this is also true for the proteins (GTPase enzymes and GTPase:effector mixtures) and incubation times intended to be used.

The model code analyses data of each GTPase assay individually and then generates pooled rate values (details on weighting of *k* for pooling and error propagation are given in S11). The analysis code allows to exclude GTPase assays from pooling if *c*_*corr*_ values or standard errors of *k* are out of range. Decision criteria are set in the code using two parameters (see steps below):

1. `conc_corr_bounds` states which range of *c*_*corr*_ allows assays to be included in the pooling. *As c*_*corr*_ *is expected to be close to 1*.*0, it is generally advisable to set in particular the lower bound not too close to 0, as such low values are indicative that the inferred activity of the GTPase is unusually low in this assay*.
2. `k_low_err_filt.’GTPase-name’` states if assays containing this particular GTPase that have low standard errors (1e-10 or lower) on the *k* values are excluded from pooling. This parameter must be declared for each GTPase individually. *In general it is advisable to set this parameter to ‘true’, i*.*e. assays with low standard errors will be excluded. If, however, a GTPase is domineered by k*_2_ *(and has k*_1_ ≈ 0 *and thus std*.*err(k*_1_)≈*0), this parameter needs to be set to ‘false’ to still allow for pooling of k. In many cases this can not be known before an initial analysis. If chosen inappropriately, the parameter simply needs to be changed and the code needs to be re-run*.

In the following we first describe the general protocol steps, which we will specify for both data of example 1 and example 2 thereafter.

#### Necessary resources

- spreadsheet editor
- Matlab software license
- Matlab analysis scripts and example data files encompassing this protocol

#### General protocol steps

1. Set up the matlab environment. To run the code, ensure that all required files are present:

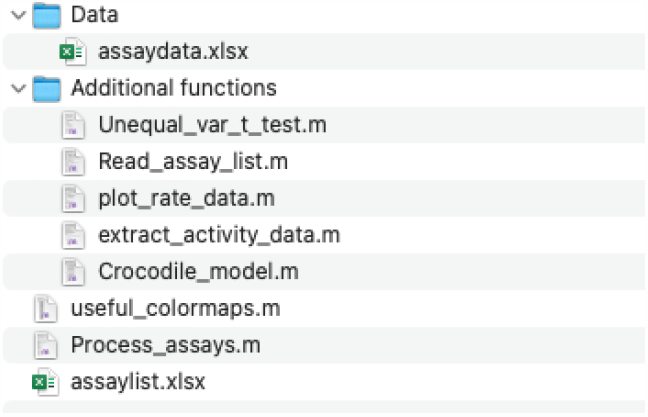 *Assay data (‘assaydata*.*xlsx’) must follow the formatting described at the end of Support Protocol 1 (Fig. 6)*. *Core of the fitting of the model occurs in the function* `Crocodile_model.m`. *This function determines which model scenario of effectors applies, and then generates fit results. To plot these*, `plot_rate_data.m` *is required, which uses colormaps from* `useful_colormaps.m` *(containing a colourmap from [Smith et al*., *2015]). When needed, an unequal variance t-test (*`Unequal_var_t_test.m`*) [Ruxton, 2006], to test for non-zero interactions between effectors. From the main function* `Process_assay.m` *a predefined spreadsheet list is read using* `Read_assay_list.m`, *to determine which data sets to use and how these are built up. The data is then read using* `extract_activity_data.m` *and passed to* `Crocodile_model.m`.
2. Define the processing of the data in the ‘assaylist.xlsx’ speadsheet. Follow the formatting provided in the example files (discussed in the next sections). The following sections are required:
  - Assay name: Gives an internal assay name. This name will be used in the output to refer to this data.
  - Proteins: These are the names of proteins whose concentration varies within the assay. State each protein in a separate column. At maximum 2 proteins can be varied per assay. GTPase ref.: State the assay name where the GTPae protein used in this assay was varied. Ensure that this assay is processed already (i.e. earlier in the list).
  - Data file location: State the relevant file location.
  - Relevant tab names: State which tab of the data file should be used.
3. Open `Process_assays.m`. Define the input parameters and processing options:

~~~
%% Input parameters
AssayListName = ‘. xlsx ’ ;
% State the a s s a y l i s t f i l e name
GTPases = { ‘’ } ;
% state proteins ( that are stated in the a s s a y l i s t f i l e ) that are GTPases
Lin Fit Effector = { ‘’ } ;
% state proteins X ( that are stated in the a s s a y l i s t f i l e ) that should be
% fitted with a linear fit : k_3, X [ GTPase ] [ X ]
Quad Fit Effector = { ‘’ } ;
% state proteins X ( that are stated in the a s s a y l i s t f i l e ) that should be
% fitted with a quadratic fit : k_3, X [ GTPase ] [ X] ^2
act_corr = true ( 1 ) ;
% Logical to determine whether terms in the rate equation that depend on
% [ GTPase ] are corrected for run− specific GTPase activity / concentration
% differences
num_draws = 1e5 ;
% Number of random draws from distribution of rate parameters k
plot _ fits = true ( 1 ) ;
% Logical indicating whether fits should be outputted in . pdf and . tiff
% ( false is faster )
print _ fits = true ( 1 ) ;
% Logical indicating whether fit results should be printed in the command window
GTP _ filt = true ( 1 ) ;
% Logical indicating whether data points 0.00 − 0.05 % GTP remaining
% should be disregarded for the fits
conc_corr_bounds = [ 0 . 5 1 . 5 ] ;
% Two−element vector that states a c_corr lower and upper bound . Assays that
% have c_corr values within t h i s range will be included in pooling of
% estimates . [ 0 Inf ] means assays with any c_corr value will be included .
% We recommend [ 0 . 5 1 . 5 ] : assays with c_corr values from 0 . 5 to 1 . 5 will be
% included .
k_low _ err _ filt . ‘GTPase−name ’ = false ;
% Logical that states per GTPase whether to discard runs from pooling that
% have very low standard errors ( 1 e−10 or lower ) on the k values .
% If nothing i s provided for a GTPase, default i s true ( i.e . runs with low
% standard errors will be excluded ) .
~~~
4. Run `Process_assays.m`. Using processing options shown above, the script will generate a folder ‘Figures’ (where all plots are saved), a spead-sheet ‘Data_summary.xlsx’ (summarising all fitting parameters) and a ‘Data_assays.mat’ file (saved in the ‘Data’ folder). ‘Data_assays.mat’ and plotting functions discussed in S10 can be used generate plots based on the fits conducted here.

#### Protocol steps: Simple assay (example 1: Ras GTPase)

We here show the specific protocol steps to run data of example 1: For this example a GTPase assay containing only serial dilutions Ras GTPase was conducted twice. Assay steps were described in the basic protocol and S2. Basic analysis of the assay data was described in Support Protocol 1 (and S9), resulting in data shown in ‘example1.xlsx’ tab: ‘matlab’. The data of each individual assay will be fitted with the model:

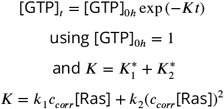

resulting one *k*_1_ and *k*2 each. A pooled estimate of *k*_1_ and *k*2 will be generated.

#### Steps

1. Set up the matlab environment. To run the code, ensure that all required files are present:

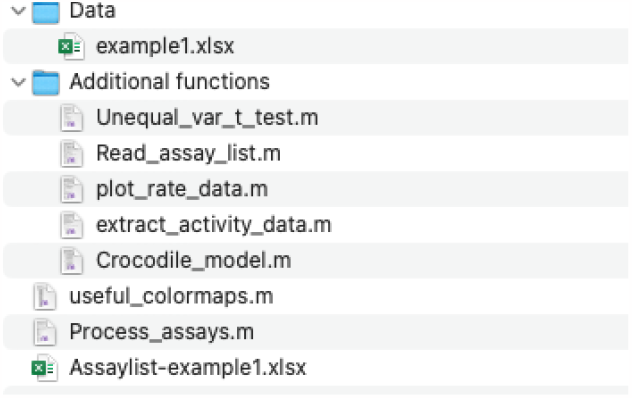
2. Define the processing of the data in the ‘assaylist-example1.xlsx’ speadsheet.

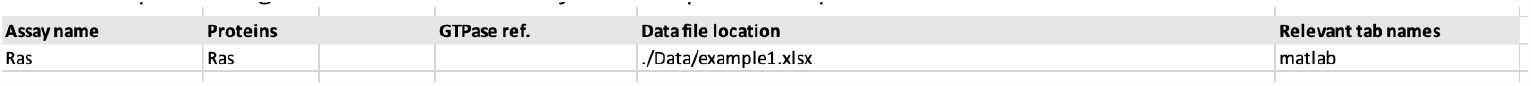 *Example 1 entails two assays of Ras serial dilutions, which are stored in ‘example1*.*xlsx’ in tab ‘matlab’. Because the concentration of Ras is varied in each assay, ‘Proteins’ states ‘Ras’*.
3. Open `Process_assays.m`. Define the input parameters:

~~~
AssayListName = ‘Assaylist −example1. xlsx ’ ;
GTPases = { ‘Ras ’ } ;
LinFitEffector = { ‘’ } ; \
Quad Fit Effector = { ‘’ } ;
act_corr = true ( 1 ) ;
num_draws = 1e5 ;
plot _ fits = true ( 1 ) ;
print _ fits = true ( 1 ) ;
GTP _ filt = true ( 1 ) ;
conc_corr_bounds = [ 0 . 5 1 . 5 ] ;
k_low _ err _ filt . Ras = false ;
~~~
4. Run `Process_assays.m`. Using processing options shown above, the script will generate a folder ‘Figures’ (where all plots are saved), a spead-sheet ‘Data_summary.xlsx’ (summarising all fitting parameters) and a ‘Data_assays.mat’ file (saved in the ‘Data’ folder). ‘Data_assays.mat’ and plotting functions discussed in S10 can be used generate plots based on the fits conducted here. The data this script ought to generate can be found in the folder ‘example1 matlab output’.

#### Protocol steps: Elaborate assay set (example 2: GTPase effector mixtures)

We here show the specific protocol steps to run data of example 2: Example 2 entails GTPase assay data (‘example2.xlsx’) of several protein mixtures (also see Fig. 2):

A. GTPase assays in which the GTPase Cdc42 is varied (all in tab: ‘dCdc42’).
B. GTPase assays using a constant Cdc42 concentration and serial dilutions of the GEF Cdc24 (all in tab: ‘Cdc42-dCdc24’).
C. GTPase assays using a constant Cdc42 concentration and serial dilutions of the GAP Rga2 (all in tab: ‘Cdc42-dRga2’). (D) GTPase assays using a constant Cdc42 concentration and serial dilutions of both GEF Cdc24 and GAP Rga2 (all in tab: ‘Cdc42-dCdc24-dRga2’).

The data of each individual assay will be fitted with the model:

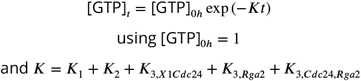

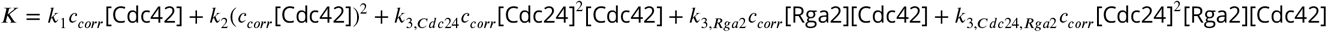

resulting for each assay one set of fitting parameters. Assays of group (A) yield *k*_1_, *k*_2_. Assays of group (B) yield *k*_3,*Cdc*24_, *c*_*corr*_. Assays of group (C) yield *k*_3,*Rga*2_, *c*_*corr*_. And assays of group (C) yield *k*_3,*Cdc*24_, *k*_3,*Rga*2_, *k*_3,*Cdc*24,*Rga*2_, *c*_*corr*_. A pooled estimate of all cycling rates *k* will be generated.

#### Steps

1. Set up the matlab environment. To run the code, ensure that all required files are present:

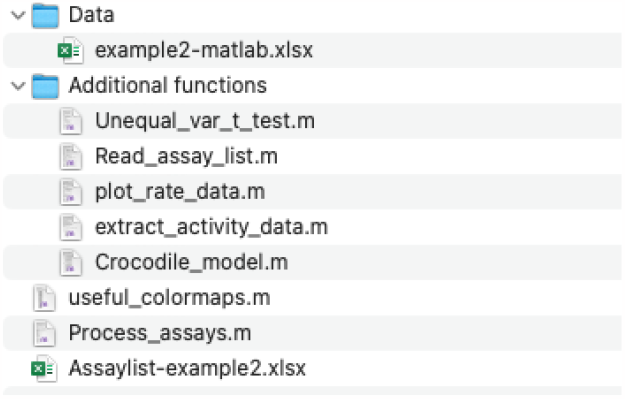
2. Define the processing of the data in the ‘assaylist-example2.xlsx’ speadsheet.

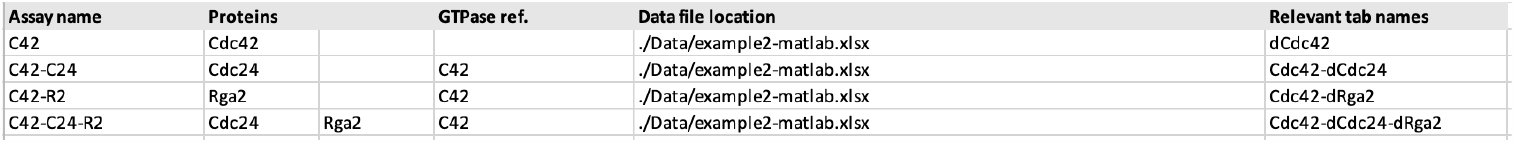 *To analyse this assay set, first all data with varied Cdc42 needs to be fitted. We give these assays the name ‘C42’. Because Cdc42 is varied, ‘Proteins’ states ‘Cdc42’*. *Then, Cdc42-effector mixtures can be analysed. We call assays with Cdc42-Cdc24 mixtures ‘C42-C24’. They include a constant concentration of Cdc42 and varied concentrations of Cdc24. Hence, ‘Proteins’ states ‘Cdc24’ (it does not state ‘Cdc42’ as well because the Cdc42 concentration is constant in these assays). To fit these assay data we want to use the rates of Cdc42 obtained in the line above, so we need to refer to ‘C42’ in ‘GTPase ref*.*’ (also see Eq. 3,4b)*. *We apply the same logic to Cdc42-Rga2 assay data, which we call ‘C42-R2’*. *We call assays in which both Cdc24 and Rga2 is varied ‘C42-C24-R2’. Here ‘Proteins’ states ‘Cdc24’ and ‘Rga2’ (because both proteins are varied), and ‘GTPase ref*.*’ states ‘C42’ to refer to the Cdc42 assay analysis in the first line*.
3. Open `Process_assays.m`. Define the input parameters:

~~~
AssayListName = ‘A s s a y l i s t −example2 . xlsx ’ ;
GTPases = { ‘Cdc42 ’ } ;
Lin Fit Effector = { ‘Rga2 ’ } ;
Quad Fit Effector = { ‘Cdc24 ’ } ;
act_corr = true ( 1 ) ;
num_draws = 1e5 ;
plot _ fits = true ( 1 ) ;
print _ fits = true ( 1 ) ;
GTP _ filt = true ( 1 ) ;
conc_corr_bounds = [ 0 . 5 1 . 5 ] ;
k_ low _ err _ filt . Cdc42 = true ;
~~~
4. Run `Process_assays.m`. Using processing options shown above, the script will generate a folder ‘Figures’ (where all plots are saved), a spead-sheet ‘Data_summary.xlsx’ (summarising all fitting parameters) and a ‘Data_assays.mat’ file (saved in the ‘Data’ folder). ‘Data_assays.mat’ and plotting functions discussed in S10 can be used generate plots based on the fits conducted here. The data this script ought to generate can be found in the folder ‘example2 matlab output’.

#### Protocol steps: Combined analysis of several assays

It is also possible to analyse assay data of example 1 and example 2 in one go. In order to do so, simply combine the inputs of the two previous examples.

1. Set up the matlab environment. To run the code, ensure that all required files are present:

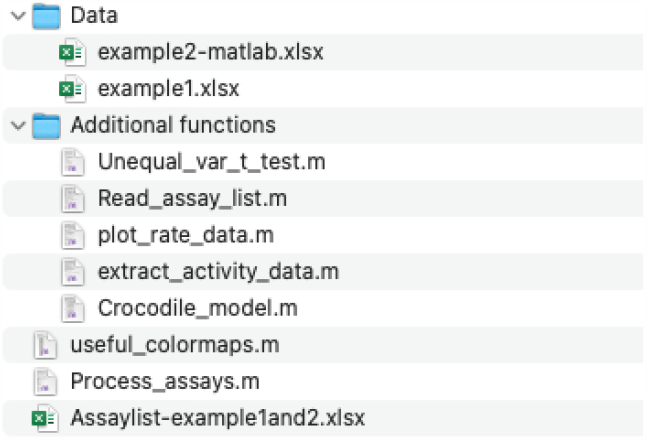
2. Define the processing of the data in the ‘assaylist-example1and2.xlsx’ speadsheet.

3. Open `Process_assays.m`. Define the input parameters:

| Assay name | Proteins |  | GTPase ref. | Data file location | Relevant tab names |
| --- | --- | --- | --- | --- | --- |
| Ras | Ras |  |  | ./Data/example1.xlsx | matlab |
| C42 | Cdc42 |  |  | ./Data/example2-matlab.xlsx | dCdc42 |
| C42-C24 | Cdc24 |  | C42 | ./Data/example2-matlab.xlsx | Cdc42-dCdc24 |
| C42-R2 | Rga2 |  | C42 | ./Data/example2-matlab.xlsx | Cdc42-dRga2 |
| C42-C24-R2 | Cdc24 | Rga2 | C42 | ./Data/example2-matlab.xlsx | Cdc42-dCdc24-dRga2 |

~~~
AssayListName = ‘Assay list −example1and2 . xlsx ’ ;
GTPases = { ‘Ras ‘, ‘Cdc42 ’ } ;
Lin Fit Effector = { ‘Rga2 ’ } ;
Quad Fit Effector = { ‘Cdc24 ’ } ;
act_corr = true ( 1 ) ;
num_draws = 1e5 ;
plot _ fits = true ( 1 ) ;
print _ fits = true ( 1 ) ;
GTP _ filt = true ( 1 ) ;
conc_corr_bounds = [ 0 . 5 1 . 5 ] ;
k_ low _ err _ filt . Ras = fa l s e ;
k _ low _ e r r _ filt . Cdc42 = true ;
~~~
4. Run `Process_assays.m`. Using processing options shown above, the script will generate a folder ‘Figures’ (where all plots are saved), a spead-sheet ‘Data_summary.xlsx’ (summarising all fitting parameters) and a ‘Data_assays.mat’ file (saved in the ‘Data’ folder). ‘Data_assays.mat’ and plotting functions discussed in S10 can be used generate plots based on the fits conducted here. The data this script ought to generate can be found in the folder ‘example1and2 matlab output’.

### Reagents and solutions

### Commentary

#### Background Information

In the GTPase assay the proteins of interest (GTPase enzymes with or without effectors) are incubated with GTP for a certain amount of time for GTPase cycling to occur, hydrolysing GTP to GDP and free phoshate. Then the reaction is stopped through addition of Glo solution, which contains a nucleoside-diphosphate kinase and ADP. The kinase converts the remaining GTP to ATP. Addition of the detection reagent, containing a luciferase/luciferin mixture, makes the ATP luminescent, which is measured using a plate reader in luminescence mode (Fig. 1) [Promega Corporation, 2015]. The luminescence signal correlates with the amount of remaining GTP (in inversly correlates with the activity of the GTPase): the higher the luminescence values, the more GTP remainded in soluion (and thus the less active the GTPase is). Low or no luminescence corresponds to very little or no remaining GTP and a higher GTPase activity.

The GTPase Glo™ assay examines the entire GTPase cycle of the respective GTPase. An assay that examines all steps of the GTPase cycle can be advantageous: For one, it allows investigation of GTPase activities and GTPase effector interactions. The obtained rates might be more comparable to those observed *in vivo* (in contrast to other *in vitro* assays that only examine one specific step of the GTPase cycle), as GTPase cycling constantly occurs *in vivo*. Further, it enables comparison of the strength of effectors that act on different steps of the GTPase cycle, such as GEFs and GAPs. Also, effector interplay can be studied. The origin of these advantages is also at the centre of the assay’s disadvantage: Because GTPase cycling (multiple completions of the entire GTPase cycle) is studied, the origin of observed effects remain elusive. It is less suitable if detailed mechanistic features of a GTPase cycle step ought to be investigated. For this, other GTPase assasy, such as the MESG/phosphorylase system [Zhang et al., 1997] or N-methylanthraniloyl-GTP/GDP [Rapali et al., 2017] system, which examine only the GTP hydrolysis or GDP release step.

#### Critical Parameters

The GTPase assay is sensitive to small pipetting errors, changes in buffer and assay components. We *strongly* recommend a mise-en-place procedure, in which each assay is carefully planned (e.g. use one of the templates provided in S1) and the materials are prepared before the assay is stared. This greatly reduce the likelihood of errors due to inattentiveness, hurry/time stress, and chaos. Accurate pipetting is essential for this assay!

Further, we advice to

- aliquot and vortex all components before use (except the Glo reagent)
- dialyse all proteins into the same buffer
- include control samples such as a buffer row in the beginning and end of the plate, a positive control (Ras GTPase enzyme), and potentially inert proteins to account for non-canonical effects
- always prepare the 2× GTP solution, 1mM ADP, and Glo solution freshly and to never re-use/store it

If executed properly, the assay leads to reliable and reproducible results (e.g. see [Tschirpke et al., 2023b, Tschirpke et al., 2023a]).

#### Troubleshooting

For a detailed troubleshooting on basic problems such as

- no change in luminescence with increasing or decreasing concentration of GTPase, GAP or GEF
- low signal-to-background ratio
- high luminescent signal
- low luminescent signal

please consult the user manual of GTPase Glo™ assay kits [Promega Corporation, 2015], we here (Tab. 4) only discuss these issues briefly.

**Table 4.**
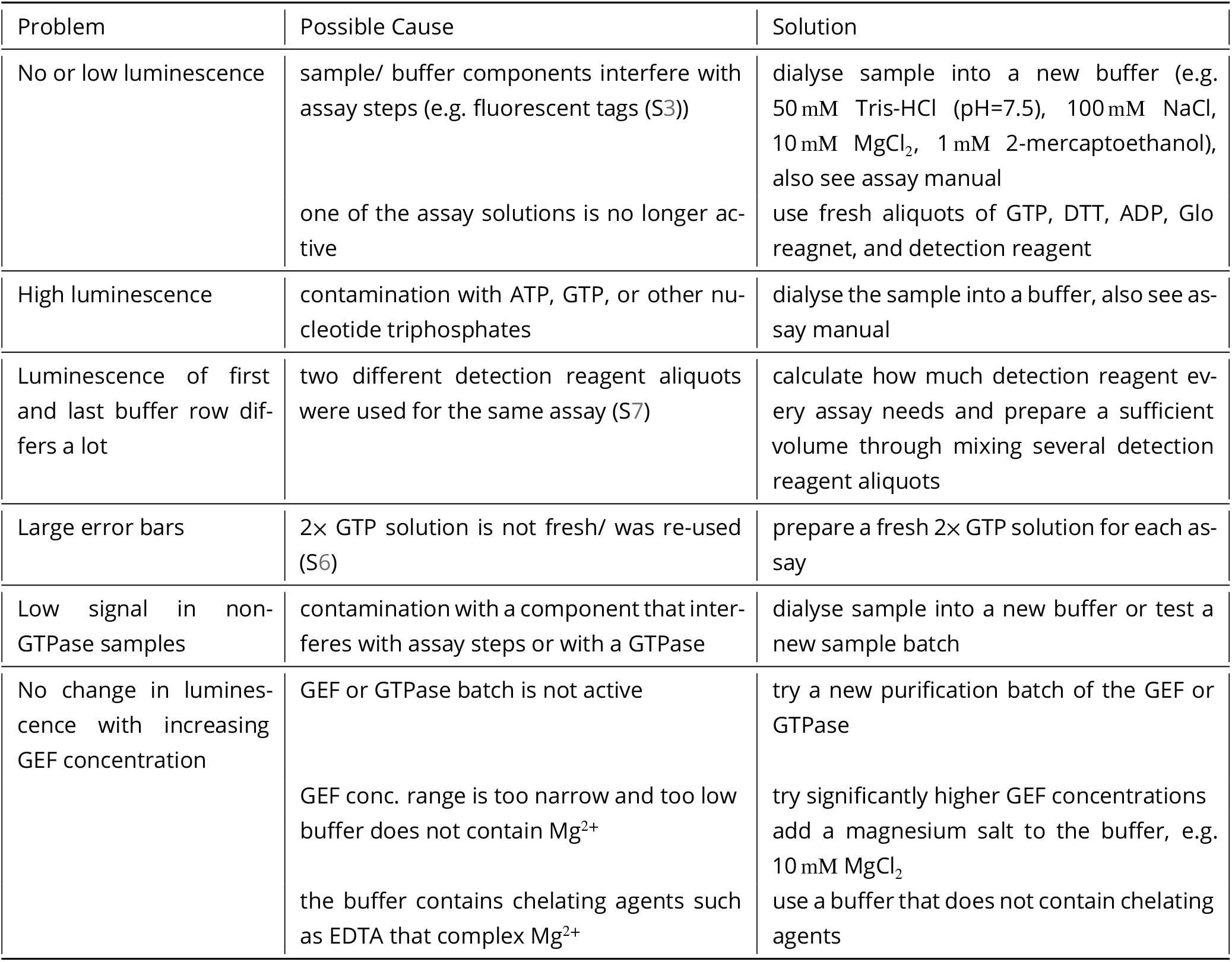
Troubleshooting guide for GTPase Glo™ assays.

| Problem | Possible Cause | Solution |
| --- | --- | --- |
| No or low luminescence | sample/ buffer components interfere with assay steps (e.g. fluorescent tags (S3))<br>one of the assay solutions is no longer active | dialyse sample into a new buffer (e.g. 50 mM Tris-HCl (pH=7.5), 100 mM NaCl, 10 mM MgCl<sub>2</sub>, 1 mM 2-mercaptoethanol), also see assay manual<br>use fresh aliquots of GTP, DTT, ADP, Glo reagent, and detection reagent |
| High luminescence | contamination with ATP, GTP, or other nucleotide triphosphates | dialyse the sample into a buffer, also see assay manual |
| Luminescence of first and last buffer row differs a lot | two different detection reagent aliquots were used for the same assay (S7) | calculate how much detection reagent every assay needs and prepare a sufficient volume through mixing several detection reagent aliquots |
| Large error bars | 2x GTP solution is not fresh/ was re-used (S6) | prepare a fresh 2x GTP solution for each assay |
| Low signal in non-GTPase samples | contamination with a component that interferes with assay steps or with a GTPase | dialyse sample into a new buffer or test a new sample batch |
| No change in luminescence with increasing GEF concentration | GEF or GTPase batch is not active<br>GEF conc. range is too narrow and too low<br>buffer does not contain Mg<sup>2+</sup><br>the buffer contains chelating agents such as EDTA that complex Mg<sup>2+</sup> | try a new purification batch of the GEF or GTPase<br>try significantly higher GEF concentrations<br>add a magnesium salt to the buffer, e.g. 10 mM MgCl<sub>2</sub><br>use a buffer that does not contain chelating agents |

#### Understanding Results

The GTPase assay yields luminescence values, which are translated into amount of remaining GTP. These values can be fitted with GTPase cycling rates *k*. Using example 1 luminescence and cycling rates *k*_1_, *k*_2_ and their pooling will be discussed. Example 2 discusses assay correction factors *c*_*corr*_ and cycling rates *k*_3,*X*_.

**Example 1**

In example 1 GTPase assay data for 5 serial dilutions of Ras GTPase were collected. The basic assay steps are described in the Basic Protocol (and S2), and their subsequent analysis is outlined in Support Protocol 1 and Support Protocol 2.

##### Luminescence

Basic protocol yields luminescence values, from which the amount of remaining GTP is calculated (Support Protocol 1). The luminescence values of the buffer rows should always be quite similar, although a small shift can be observed (Fig. 7 (top)). If this shifts gets big, the error bars of the entire assay will increase (because the buffer wells are used for normalisation). Dilutions of Ras need to show lower luminescence values than the buffer (because Ras is a GTPase and hydrolyses GTP, which correlates with luminescence (Fig. 1)). With increasing Ras concentration, the luminescence should drop. Replicas/wells containing the same Ras concentration should have the same luminescence (Fig. 7 (bottom)). The bigger their spread, the bigger the error will be. In some cases we observe a small decrease of the luminescence over time (Fig. 7). As long as this decrease is not drastic and occurs in all wells to roughly the same extend, it will not impact the analysis. Assays in which the luminescence decreases drastically within 20 min should be discarded from further analysis, as well as wells that behave differently than the majority.

**Figure 7.**
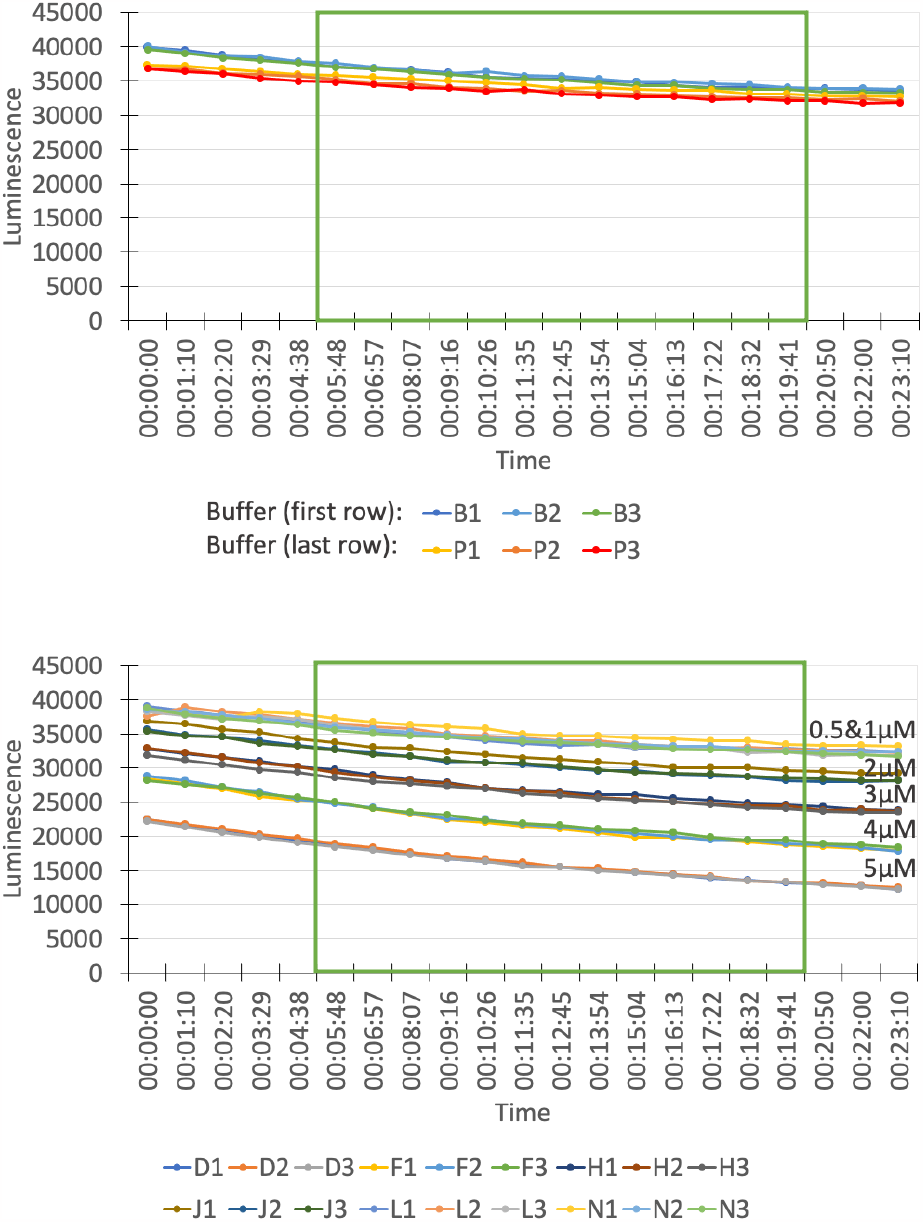
Example 1: Measured luminescence values of buffer wells (top) and Ras GTPase dilutions (bottom) over the time course of the measurement. The green box indicates values that are used for averaging.

According to the assay manual, the luminescence should be measured after 5 min. We average the luminescence values from 5-20 min for every assay, to smoothen out possible small deviations in the luminescence curve. The exact time window (as long as it begins after 5 min) is less important than choosing exactly the same time window for every assay.

The luminescence values are used to calculate the amount of remaining GTP (Fig. 5). It is possible to use these values to compare the activity of different GTPases, the activity of two different effectors on a GTPase, or the activity of one effector on different GTPases, as long as all samples are part of the same assay (to avoid small variability between assays) and are used in the same concentration.

##### GTPase cycling rates *k*_1_, *k*_2_

To increase comparability of GTPase and effector activities, we developed a GTPase cycling model (S8). Support Protocol 2 describes how the model can be used to fit data of Ras. In short, we fit an exponential to the amount of remaining GTP (S8):

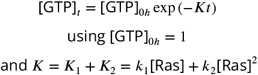

Here *K* refers to the ‘overall GTP hydrolysis rate’ and depends on the concentration of the GTPase Ras. GTPase cycling rates *k* describe the rate of the *entire* GTPase cycle of Ras.

The model fits two cycling rates (*k*_1_ and *k*_2_) for every GTPase (Eq. 3). *K*_1_ accounts for the linear contribution of the GTPase (*K*_1_ = *k*_1_ [GTPase]). *K*_2_ accounts for the quadratic contribution (*K*_2_ = *k*_2_ [GTPase]^2^) which can be due to dimerisation or other cooperative effects increasing the activity of the enzyme (for examples, see [Zhang et al., 1999]). The bigger the *K*_2_/*K*_1_ ratio, the bigger the contribution of the non-linear term. This can suggest that the GTPase forms a dimer that is more active than the monomer. However, additional studies examining GTPase di-/oligomerisation are required before making such a deduction. The data indicates that some effect leads to a non-linear concentration dependence of the rate *K*. It does not reveal its origin. Ras does not show a linear concentration dependence (Fig. 8) and is domineered by *k*_2_ (Fig. 9: *k*_2_ >>> *k*_1_). In this case this is likely due to dimerisation - Ras GTPases are known to dimerise.

**Figure 8.**
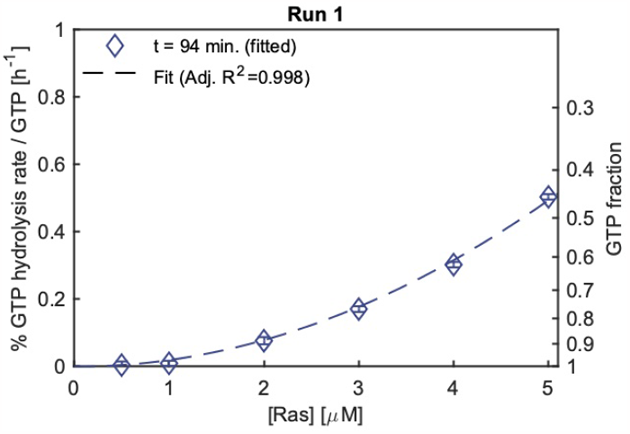
Example 1: The overall GTP hydrolysis rate *K* does not scale linearly with Ras concentration. (The figure is automatically generated when running the matlab code (using ‘plot_fits = true(1)’).)

**Figure 9.**
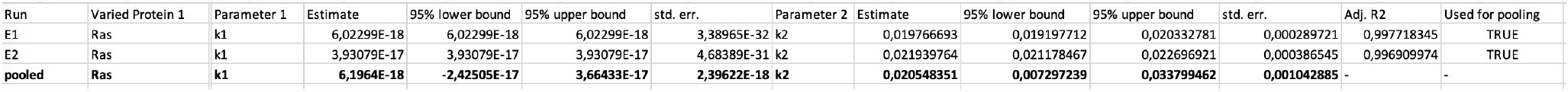
Example 1: GTPase cycling rates *k*_1_ and *k*_2_ for Ras.

##### Output file structure and pooling of rates *k*

Estimates of Ras GTPase cycling rates *k*_1_ and *k*_2_ are summarised in ‘Data_summay.xlsx’ (Fig. 9). The file is structured in the following way: The column ‘Run’ shows the assay number, followed by two ‘varied protein’ colmuns stating which protein(s) were varied in the assay (i.e. for which proteins several dilutions were used). If the assay contains only one varied protein, the second column is empty (as it is the case for example 1: here only Ras was used). Then Parameter 1 is named, and its values, 95% upper and lower bound, and standard error is shown. The same is repeated for possible parameters 2 and 3. The last column show adjusted R^2^ of the fit and a Boolean if the experiment is used for pooling (‘Used for pooling’).

For each assay (row, ‘Run’: E1, E2) the rate value estimates are shown. Below, in the row which contains ‘pooled’ in the ‘Run’ column, the pooled values of *k* estimates are stated . The column ‘varied protein’ states which proteins were varied in the assay (i.e. for which proteins several dilutions were used). ‘Used for pooling’ indicates if experiment is used for pooling. An experiment can be excluded from pooling if *c*_*corr*_ or standard errors of *k* are out of range. Decision criteria are set before analysing the data using two parameters (see Support Protocol 2): (1) `conc_corr_bounds` states which range of *c*_*corr*_ allows assays to be included in the pooling. (2) `k_low_err_filt.Ras` states if assays with low standard errors (1e-10 or lower) on the *k* values are excluded from pooling. In Support Protocol 2 (example 1) we used

~~~
conc_corr_bounds = [ 0 . 5 1 . 5 ] ;
k _ low _ err _ filt . Ras = false ;
~~~

for analysing Ras data. `conc_corr_bounds` states which range of *c*_*corr*_ allows assays to be included in the pooling. It does not influence pooling of *k*_1_ and *k*_2_, as *c*_*corr*_ only is used for fitting *k*_3,*X*_ (Eq. 4,5). We set `k_low_err_filt.Ras` to ‘false’ - experiments with low standard errors on *k* values are *not* excluded from pooling. This makes sense for Ras, as Ras is dominated by *k*_2_ has *k*_1_ ≈ 0 (and thus *k*_1_ errors are close to 0).

**Example 2**

For this example data of a set of GTPase assays is provided, including GTPase assays of

A. Cdc42 dilutions (GTPase enzyme),
B. Cdc24 (GEF) dilutions using a constant Cdc42 concentration,
C. Rga2 (GAP) dilutions using a constant Cdc42 concentration, and
D. Cdc24 (GEF) dilutions and Rga2 (GAP) dilutions using a constant Cdc42 concentration.

The analysis is described in Support Protocol 2. In short, a GTPase cycling model (S8) is fitted:

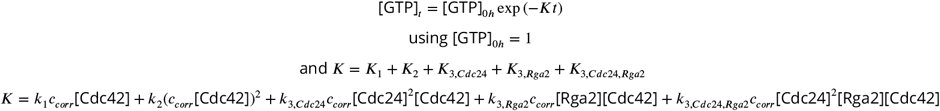

Here *K* refers to the ‘overall GTP hydrolysis rate’ and depends on the concentration of the GTPase (and effector *X*). First data of Cdc42 serial dilutions (data (A)) are fitted with an exponential to obtain estimates of GTPase cycling rates *k*_1_ and *k*_2_ for each experiment. These estimates are then bundled to get a isngle pooled estimate for *k*_1_ and *k*_2_. Then, Cdc42 - Cdc24, Cdc42 - Rga2, and Cdc42 - Cdc24 - Rga2 mixtures (data (B), (C), (D)) are fitted to obtain estimates for the correction factors *c*_*corr*_ and cycling rates *k*_3_ (measuring the impact of the effector on GTPase cycling) for every experiment. These latter estimates can then be pooled to get single estimate for the *k*_3_’s (a pooled estimate for *c*_*corr*_ has no function here).

##### Correction factors *c*_*corr*_

The correction factor *c*_*corr*_ accounts for variability between assays, i.e. for the observation that the rates for the GTPase can vary between assays. Possible reasons for this include small concentration differences introduced though pipetting of small volumes (as are required for this assay), temperature and shaker speed fluctuations during the incubation step, small changes in effective GTPase concentration through sequestration of GTPases in complexes with effectors, and/or intrinsic changes in the protein activities due to other external conditions. *c*_*corr*_ maps all factors that lead to variations between assays onto the GTPase concentration. The correction factor is estimated for each experiment individually, relative to a predefined GTPase serial dilution assay. This is done by matching the overall hydrolysis rate at zero effector concentration inferred from the fit, to the estimated overall hydrolysis rate based on the GTPase serial dilution assay. Concretely, *c*_*corr*_ follows from:

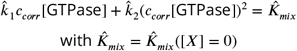

Where 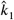 and 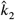 are the estimates of cycling rates from the GTPase serial dilution assay and 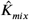 are the estimated median of the overall hydrolysis rate fit at zero effector concentration value (i.e. 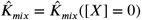). By using random draws of the fitted overall hydrolysis rate distribution, we can also obtain a distribution for *c*_*corr*_ as well. Consequently, *c*_*corr*_ = 1.0 means that relative to the serial dilution assay with the same GTPase (= reference assay), the GTPase in this assay has the same intrinsic activity, which is optimal. This is optimal. *c*_*corr*_ = 0.5 means that relative to the reference assay, the GTPase in this assay is 50% less active, and *c*_*corr*_ = 1.5 means that that the GTPase in this assay is 50% more active. Very low correction values indicate that in the assay the GTPase is behaving differently than before (possibly because the conditions are different) or that a fundamental error occurred (e.g. large pipetting error). To ensure consistency within an assay set we advice to disregard that assay from further analysis.

To implement this recommendation, we used `conc_corr_bounds = [0.5 1.5]` for analysing the data set in Support Protocol 2 (example 1). This means that assays yielding *c*_*corr*_ values lower than 0.5 and larger than 1.5 will be excluded from pooling. Fig. 10 shows a histogram of all correction factors. Apart from one assay, which yielded *c*_*corr*_ ≈ 0.1, all *c*_*corr*_ values are within range, with most values being close to 1.

**Figure 10.**
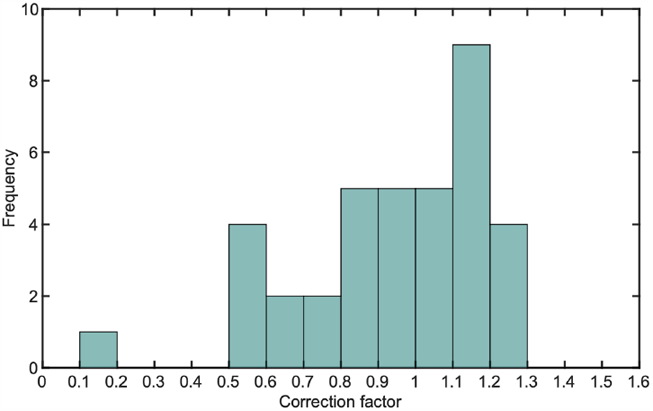
Example 2: Histogram plot of *c*_*corr*_ using a bin size of 0.1. Generated for all *c*_*corr*_ values of example 2 (using ‘Plot_c_corr_histogram.m’ (S10) and ‘example2-histogram.xlsx’ in the folder ‘example2 matlab output’).

‘Data_summay.xlsx’ states correction factors as ‘parameter 1’ for assays in which either Cdc24 or Rga2 or both effectors are varied (‘Varied protein 1’, ‘varied protein 2’) Correction factor errors are discussed in S11.

##### GTPase cycling rates *k*_3_

GTPase cycling rates *k* describe the rate of the *entire* GTPase cycle of the GTPase, or the effect of an effector on the overall rate. Cycling rates *k*_3,*X*_ can thus describe how strongly an effector *X* affects the GTPase cycle, but do not reveal which step of the GTPase cycle the effector is acting upon or how strongly the effector is effecting this step (or if the effector acts on multiple steps). To investigate these mechanism, other GTPase assays (e.g. those mentioned in the background information section), need to be conducted.

Assays of group (B) are used to fit cycling rates *k*_3,*Cdc*24_ which describe the effect of the GEF Cdc24 on Cdc42’s overall GTPase cycle. We choose a quadratic fit (*K*_3,*Cdc*24_ = *k*_3,*Cdc*24_[*Cdc*42][*Cdc*24]^2^) because the data showed a non-linear dependence of the overall rate *K* on the Cdc24 concentration (Fig. 11). The fit is a phenomenological description of the data and does not reveal where the non-linearity originates from. One possibility could be dimerisation, with Cdc24 dimers exhibiting an increased activity [Tschirpke et al., 2023a]. *k*_3,*Cdc*24_ values are shown ‘Data_summay.xlsx’ as ‘parameter 2’ for assays in which Cdc24 is varied (‘Varied protein 1’).

**Figure 11.**
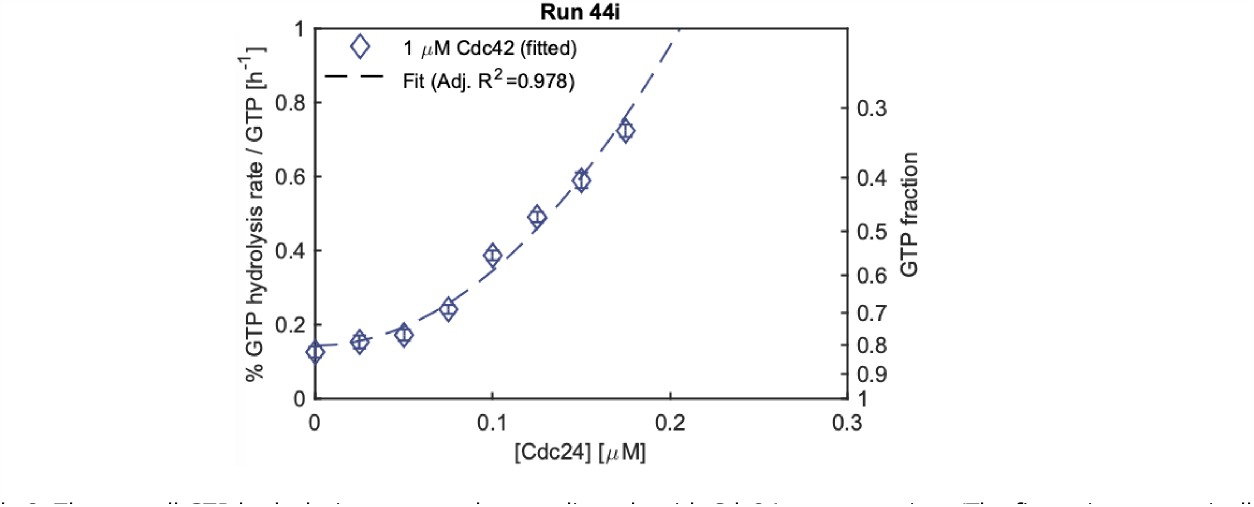
Example 2: The overall GTP hydrolysis rate *K* scales non-linearly with Cdc24 concentration. (The figure is automatically generated when running the matlab code (using ‘plot_fits = true(1)’).)

In a similar fashion, assays of group (C) are used to fit cycling rates *k*_3,*Rga*2_ which describe the effect of the GAP Rga2 on Cdc42’s overall GTPase cycle. A linear fit (*K*_3,*Rga*2_ = *k*_3,*Rga*2_[*Cdc*42][*Rga*2]) was used to describe the data.

Data of group (D) contains Cdc42 - Cdc24, Cdc42 - Rga2, and Cdc42 - Cdc24 - Rga2 mixtures. In these assays both Cdc24 and Rga2 are varied (‘varied protein 1’, ‘varied protein 2’). The data is fitted with 3 rate parameters (Fig. 12): *k*_3,*Cdc*24_ (= ‘parameter 2’), *k*_3,*Rga*2_ (= ‘parameter 3’), *k*_3,*Cdc*24,*Rga*2_ (= ‘parameter 4’). *k*_3,*Cdc*24_ describes the effect of Cdc24 on the entire GTPase cycle and *k*_3,*Rga*2_ describes the effect of Rga2. These rates should be close to those obtained in assays including only Cdc42 - Cdc24 (or Cdc42- Rga2 mixtures) (Fig. 13). a large difference can indicate a problem with the fit or the assay. The cycling rate *k*_3,*Cdc*24,*Rga*2_ accounts for any potential synergy between both proteins. *k*_3,*Cdc*24,*Rga*2_ = 0 means that there is no interaction between both proteins. *k*_3,*Cdc*24,*Rga*2_ < 0 suggests that the proteins antagonise/ inhibit each other. *k*_3,*Cdc*24,*Rga*2_ > 0 suggests that there is synergy. The term states if there is an interaction between both protein, i.e. if the combined effect of both proteins is bigger than the sum of their individual contributions to the *overall* rate. (The significance of the deviation from zero is is also tested using [Ruxton, 2006]. The p-value for the null hypothesis of zero interaction is stored in the ‘Data_assays.mat’ output file in the ‘Assays_processed’ variable, in the field of that assay, in the sub-field ‘k_tot’, under ‘Interaction_p’.) While a positive *k*_3,*Cdc*24,*Rga*2_ indicates that there is some synergy when both Cdc24 and Rga2 are added to Cdc42, it does not reveal the origin of the synergy. The origin could be due to physical protein-protein interactions or due to rate-limiting steps in the cycle, which are relieved once both proteins are added [Tschirpke et al., 2023a]. One has to thus be careful in the interpretation of this rate.

**Figure 12.**
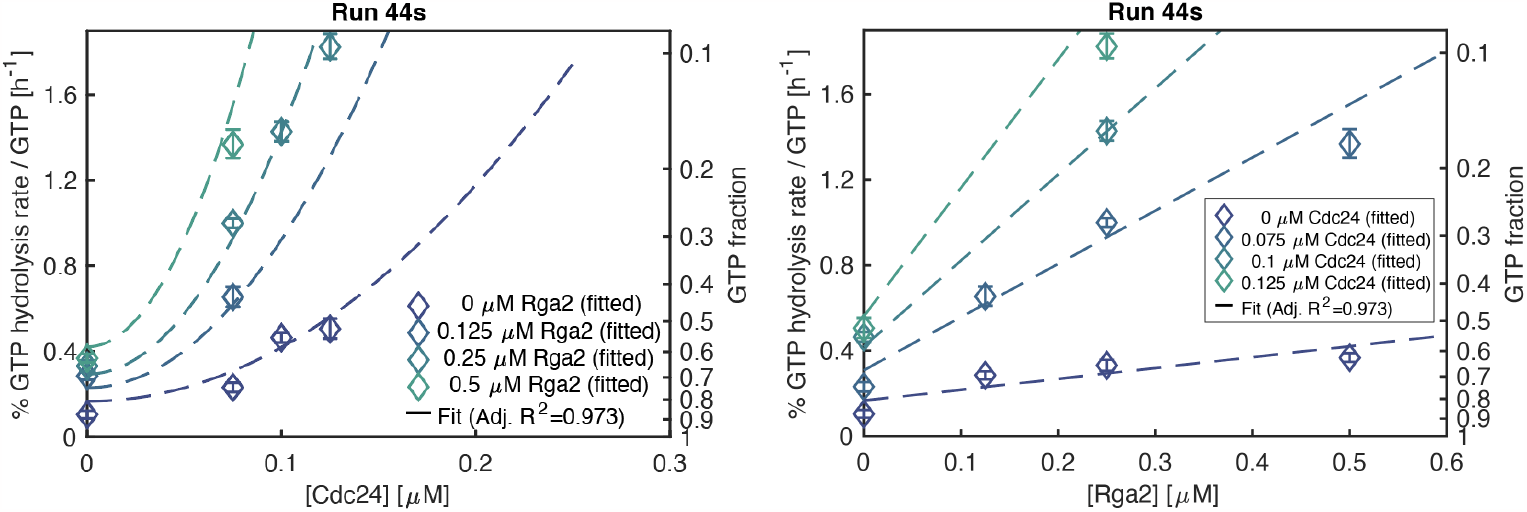
Example 2: Fit of Cdc42-Cdc24-Rga2 mixtures. (The figures are automatically generated when running the matlab code (using ‘plot_fits = true(1)’).)

**Figure 13.**
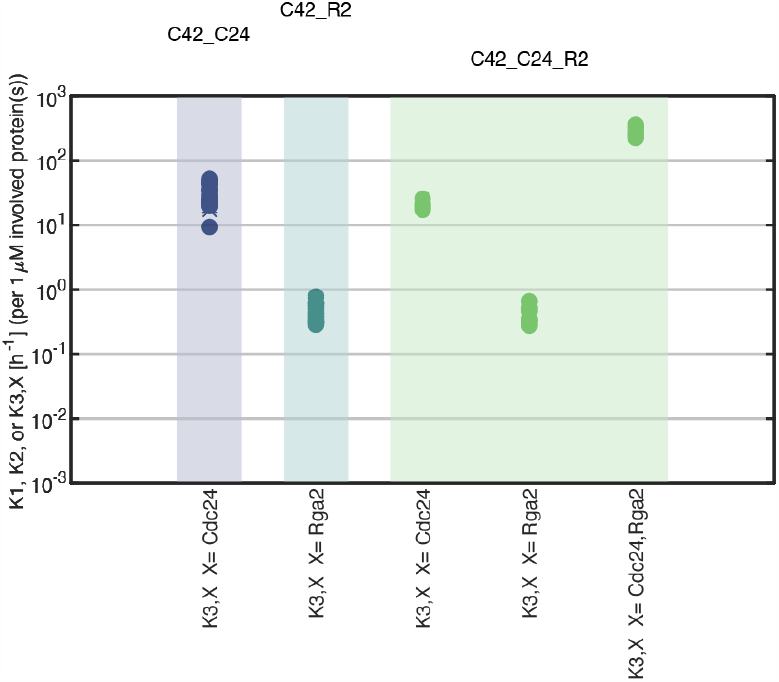
Example 2: Plot of rates *K*_3_, *X* for Cdc42-Cdc24 (left: ‘C42-C24’), Cdc42-Rga2 (middle: ‘C42-R2’), and Cdc42-Cdc24-Rga2 (right: ‘C42-C24-R2’) assays. Rate values of individual experiments are shown as filled dots. The average is shown as a cross and error bars represent the standard error. Generated for assay names ‘C42-C24’, ‘C42-R2’, and ‘C42-C24-R2’ (using ‘Plot_pooled_values_plot_std_err.m’ (S10) and ‘Data_assays.mat’ in the folder ‘example2 matlab output’).

##### Comparison of cycling rates *k*

Overall GTP hydrolysis rates *K* are concentration dependent and have the unit *h*^−1^. Cycling rates *k* are concentration-independent, but have different units and are therefor difficult to compare: *k*_1_ : [µM^−1^*h*^−1^], *k*_2_ : [µM^−2^*h*^−1^], *k*_3,*Cdc*24_ : [µM^−3^*h*^−1^], *k*_3,*Rga*2_ : [µM^−2^*h*^−1^], *k*_3,*Cdc*24,*Rga*2_ : [µM^−4^*h*^−1^]. The easiest way to represent all rates in one plot is to plot overall GTP hydrolysis rates *K* for 1 µM of each protein (Fig. 13). However, one has to consider the different concentration-dependencies when interpreting/comparing the rates *K*. A rate that scales quadratically with protein concentration will be twice as big as a rate that scales linearly when doubling the protein concentration. Some rates might also just be valid for the regime in which they were fitted in (they might show saturation in higher concentration regimes). Thus, one has consider which protein concentrations one uses to calculate *K* (in order to make cycling rates *k* comparable).

#### Time Considerations

GTPase assays usually take a few hours, of which time is consumed by incubation steps. For an assay involving 8 samples, including one Ras postive control and 5 serial dilutions of the GTPase of interest (Fig. 3), using an incubation time of 1.5 h, we estimate 0.5 h for preparation of materials and solutions, 2 h for conducting the assay, and another 0.5 h for the luminescence readout. This amounts to a total of 3 h, of which 2 h account for incubation/measurement steps.

Assays encompassing more samples may involve longer preparation times (as more protein dilutions need to be prepared), and longer incubation times subsequently also prolong the total amount of time spent.

If proteins need to be dialysed into a suitable buffer first, an additional overnight step of dialysis is required.

The analysis time depends on the level of automatisation and can range from only a few to 30-40 min. If the first step is done manually in Excel, we estimate 30 min for this step. The following analysis in Matlab requires only a few minutes of runtime.

## Data Sharing and Data Availability

The data that support and exemplify the protocols are openly available at https://data.4tu.nl at http://doi.org/10.4121/ac196f25-1c20-4c0c-a0b9-f01cd3fadc45.

## Acknowledgements

We thank D. McCusker (University of Bordeaux) for the plasmid pDM272 and N. Dekker (TU Delft) for the plasmid pET28a-His-mcm10-Sortase-Flag.

L. Laan gratefully acknowledges funding from the European Research Council under the European Union’s Horizon 2020 research and innovation programme (grant agreement 758132) and funding from the Netherlands Organization for Scientific Research (Nederlandse Organisatie voor Wetenschappelijk Onderzoek) through a Vidi grant (016.Vidi.171.060).

## Contributions

**S. Tschirpke:** Conceptualization, Methodology, Validation, Investigation, Formal analysis, Writing - Original Draft, Writing - Review & Editing, Visualization, Project administration. **W. K.-G. Daalman:** Conceptualization, Methodology, Validation, Investigation, Formal analysis, Writing - Original Draft, Writing - Review & Editing, Software. **L. Laan:** Writing - Review & Editing, Project administration, Funding acquisition, Supervision.

## Supplement S1

The following pages include preparation sheets for conducting GTPase assays with 1, 2, or 3 sample rows.

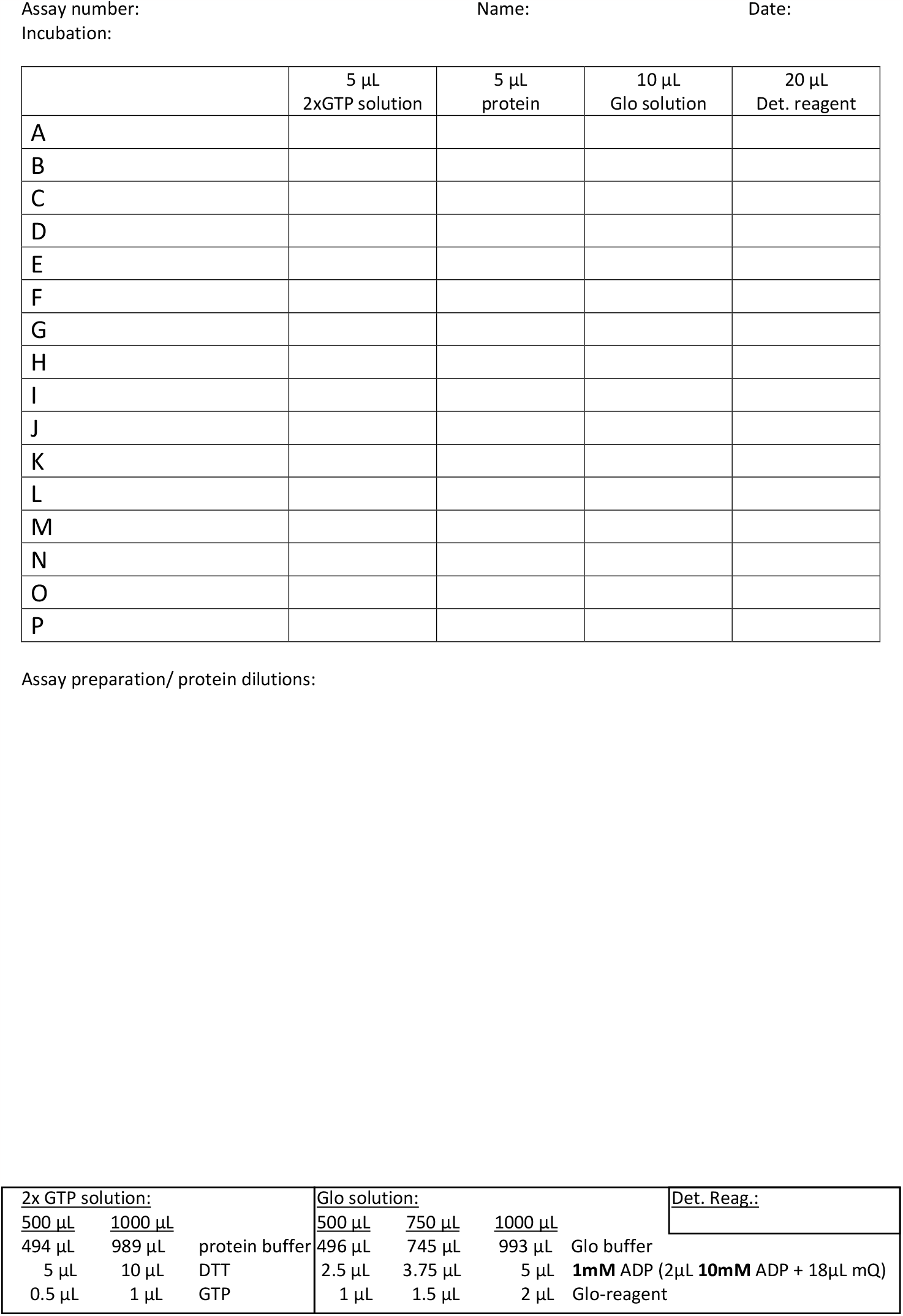

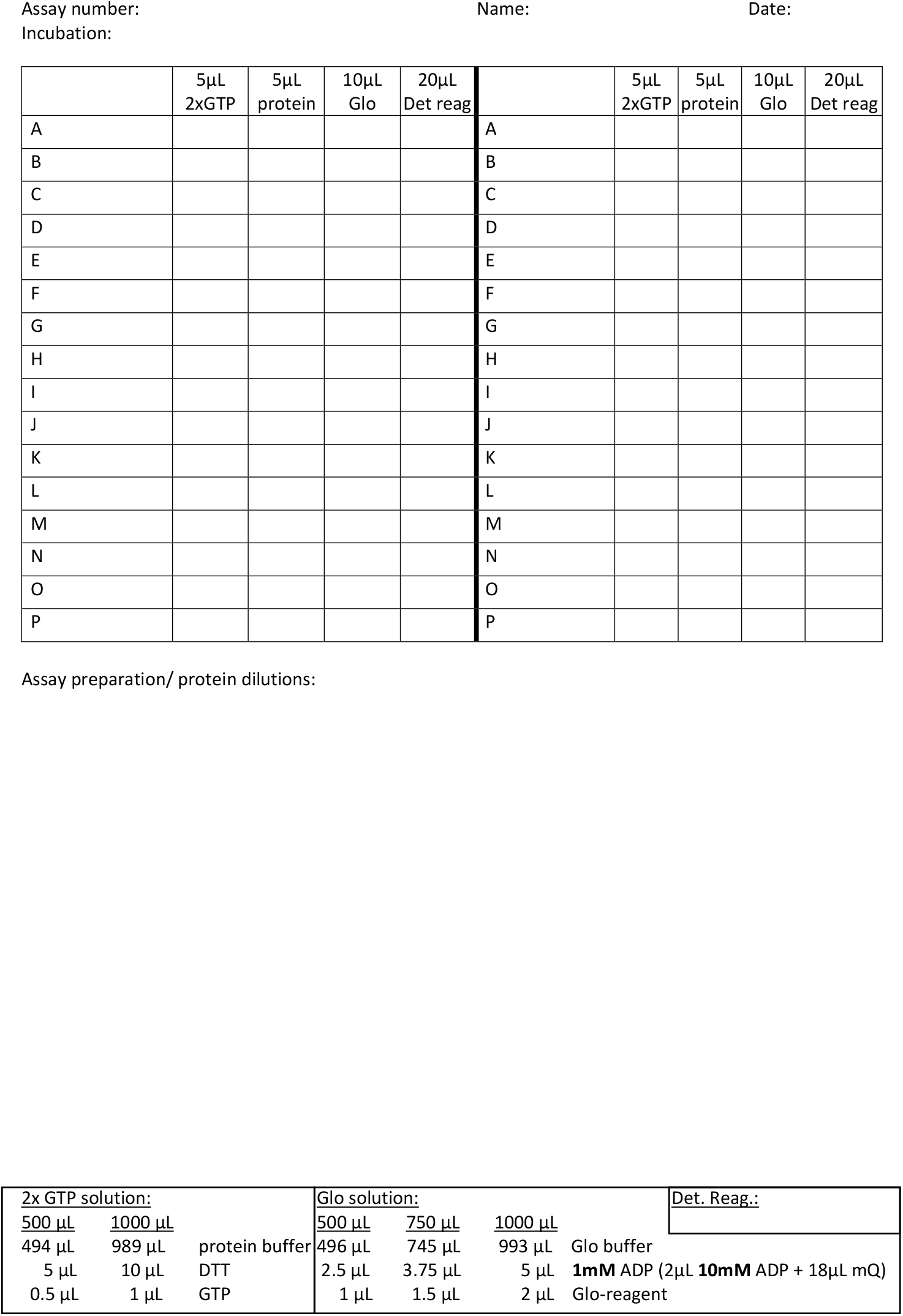

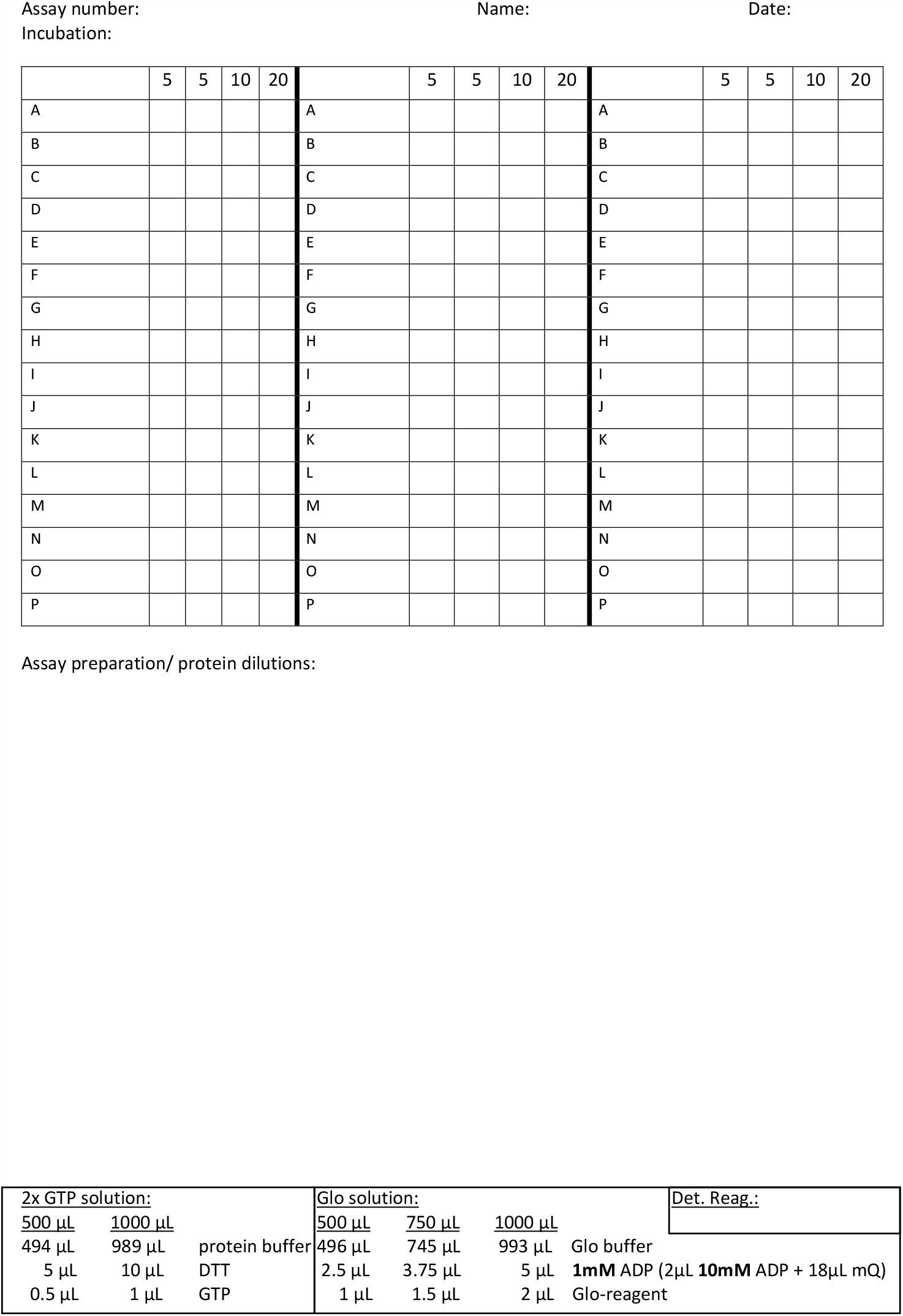

### Supplement S2

This supplement shows a prepared plate and preparatory sheet for a GTPase assay of 6 Ras GTPase serial dilutions (example 1) (see basic protocol). It leads to data that can be found in ‘example1.xlsx’, tab: ‘E1’, ‘E2’.

**S2 Figure 1.**
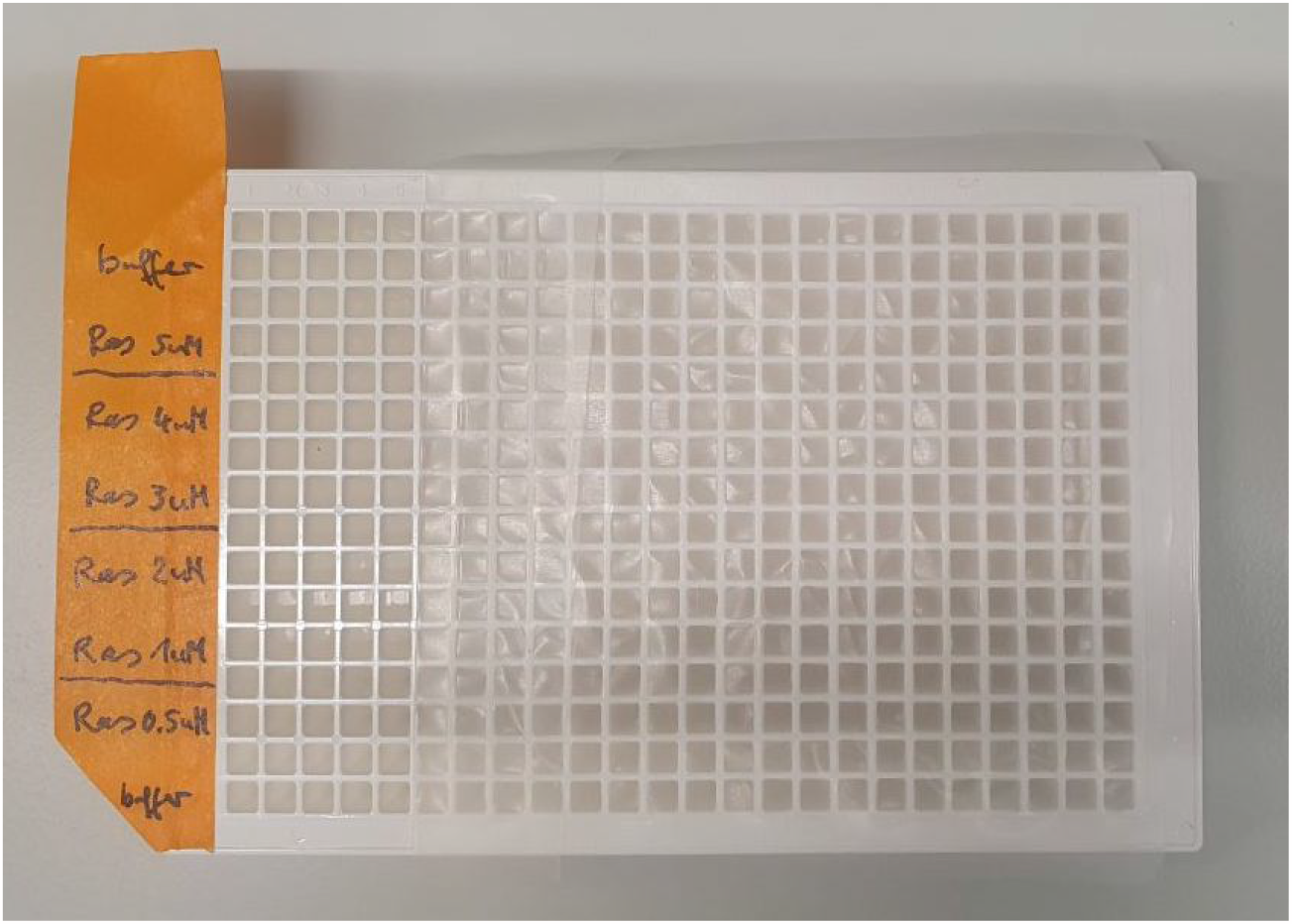
Preparation of a plate for a GTPase assay containing 6 Ras GTPase serial dilutions (example 1).

**S2 Figure 2.**
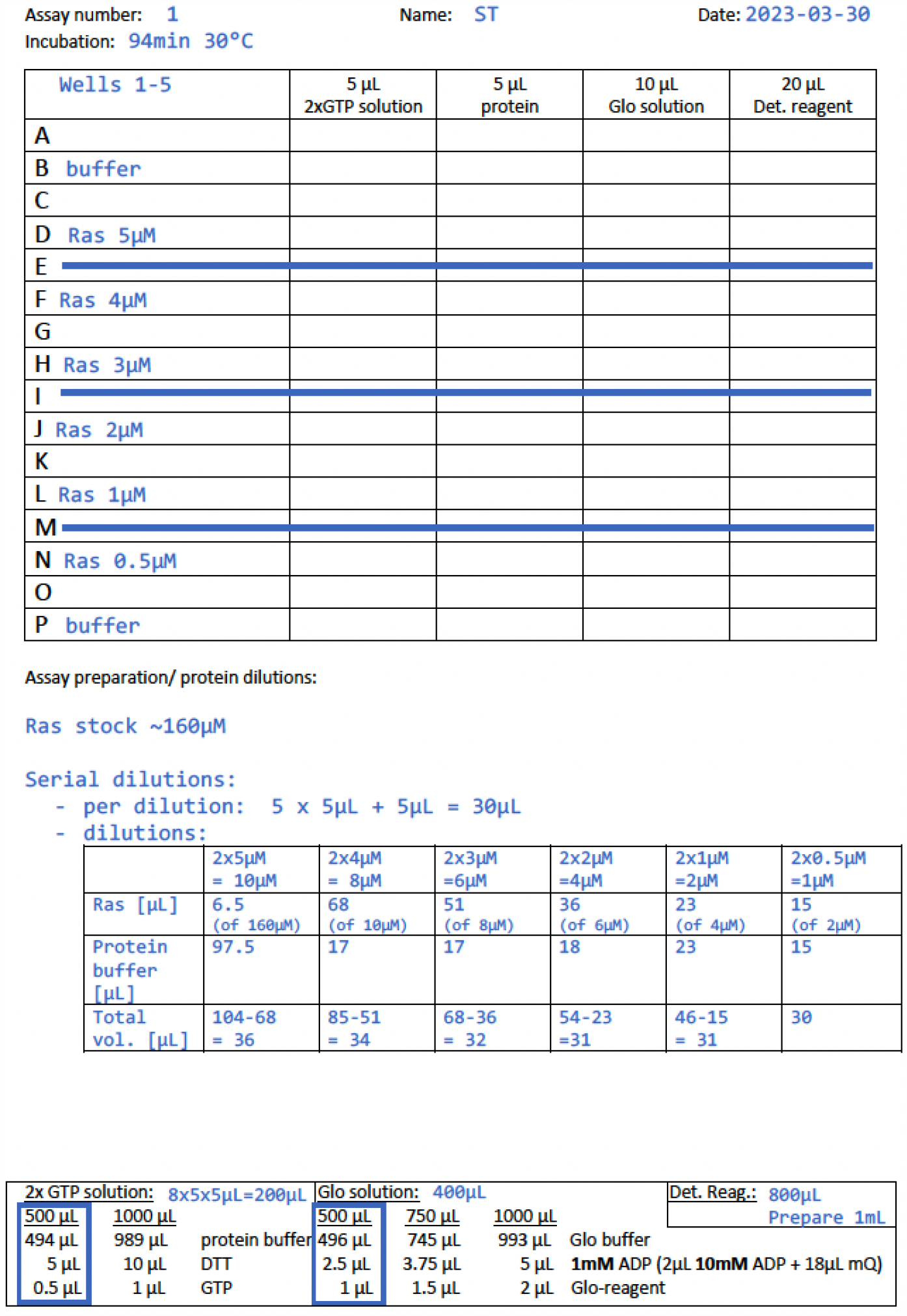
Preparation of a GTPase assay of 6 Ras GTPase serial dilutions (example 1) (made using templates provided in S1).

### Supplement S3

Before including fluorescently labelled proteins into the GTPase assay, assess if the fluorescent tags interfere with the assay readout (= luminescence). Some fluorescent tags (e.g. Alexa488) reduce the luminescence almost completely (e.g. mNeon-green), while others only affect it mildly (e.g. Alexa488) (S3 Fig. 1).

**S3 Figure 1.**
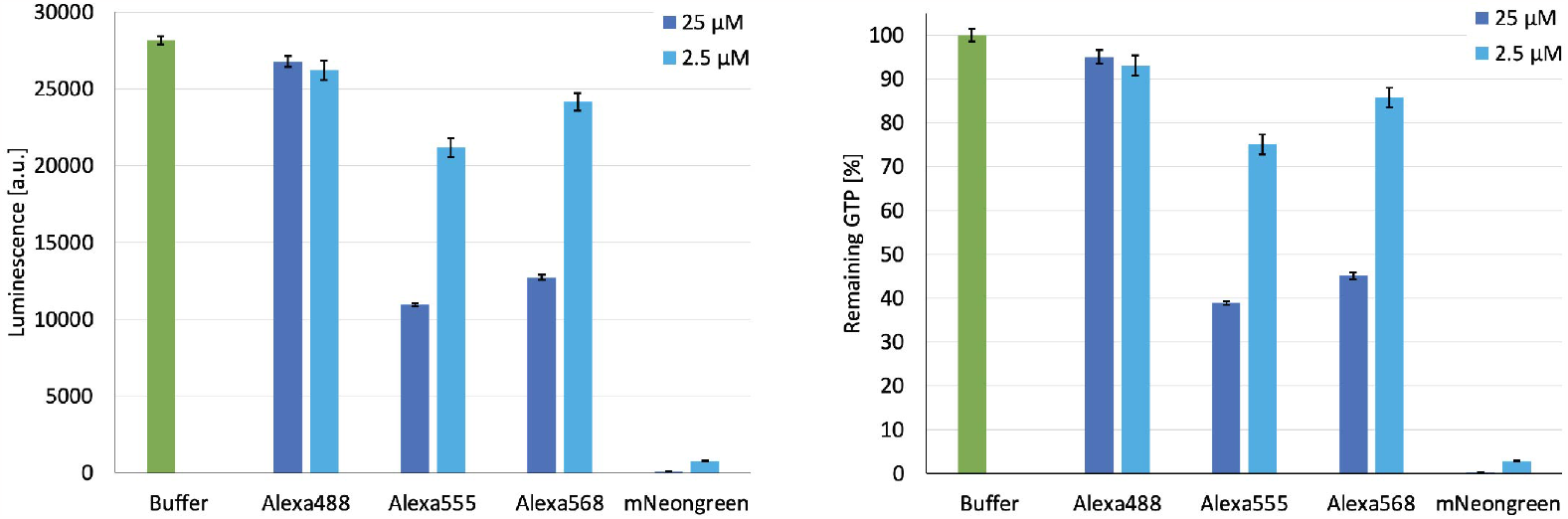
Fluorescent tags can interfere with the GTPase assay readout: Luminescence (left) and perceived amount of remaining GTP (right) of four fluorophores (normalised to buffer). Fluorophores are not GTPase enzymes and are not expected to hydrolyse GTP. A drop in luminescence signal (and thus a decrease in the perceived amount of remaining GTP) is likely due to absorption of some of the luminescence signal by the fluorophores.

### Supplement S4

**S4 Figure 1.**
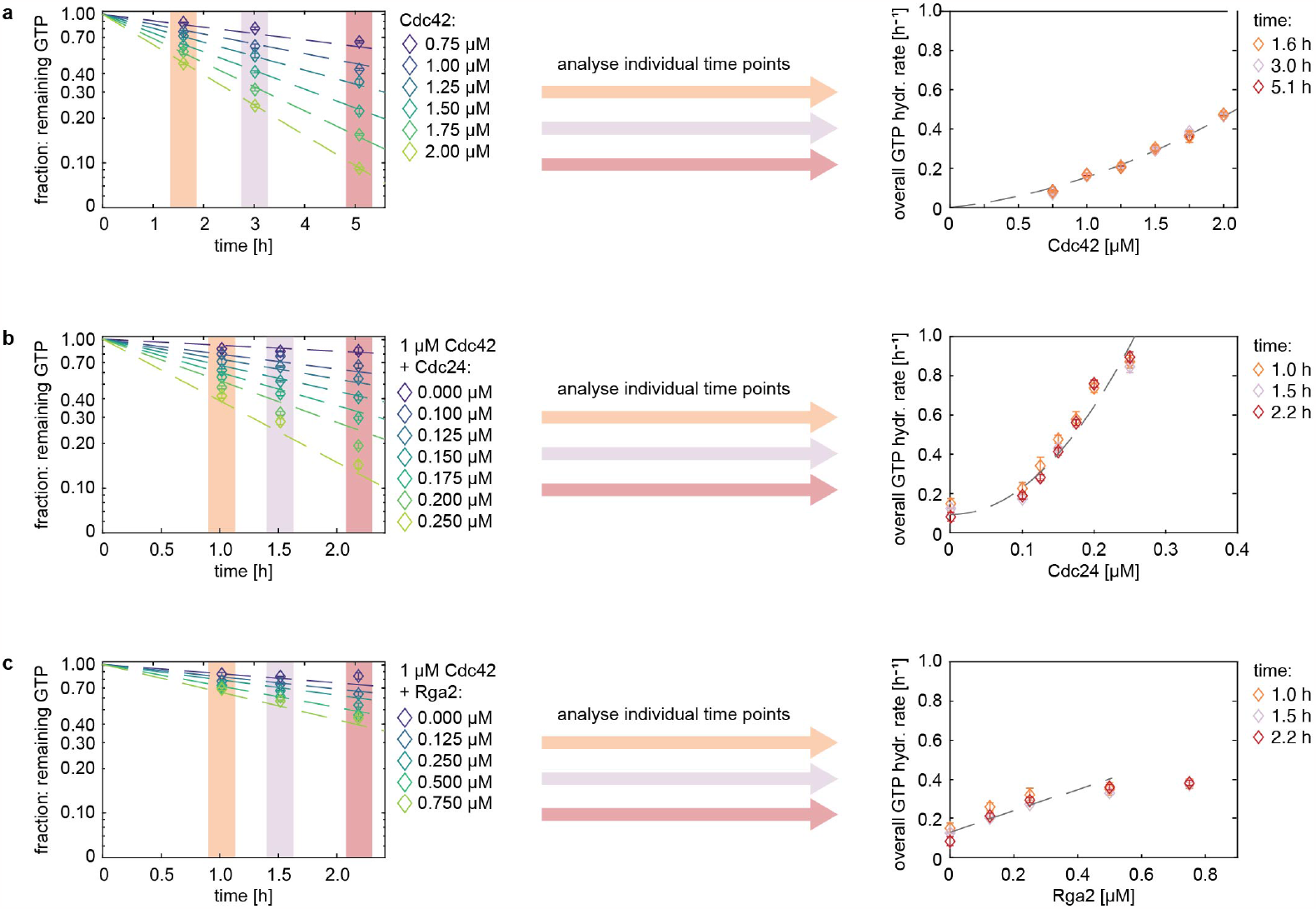
The GTP concentration declines exponentially with time in GTPase reactions. Amount of remaining GTP for (a) Cdc42 concentrations (b) Cdc42 Cdc24 mixtures, and (c) Cdc42 Rga2 mixtures, each for three time points (measured as one individual assay per time point). The remaining GTP content declines exponentially with time (left). Data of each individual time point shows the same overall GTP hydrolysis cycling rate for each GTPase - effector mixture. Thus, only one time point per assay condition is needed, to fit the data (right).

### Supplement S5

**S5 Figure 1.**
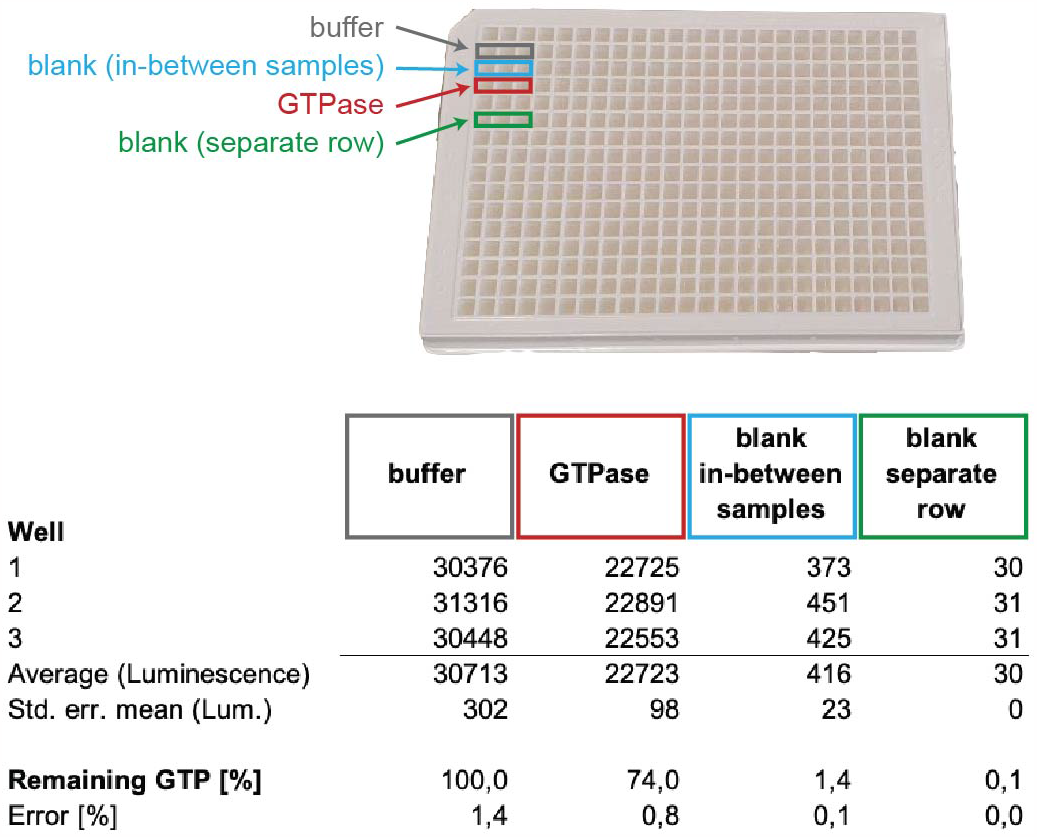
Leave one empty row between all sample rows to avoid any spill-over of luminescence signal between samples. The row ‘blank (in-between samples)’ (light blue) does not contain any solution, it is empty. It is placed between a row that contains buffer and a row with a GTPase sample, both of which have a strong luminescence signal (as is expected). This leads to a small, but detectable luminescence signal in this row. In comparison, a similar blank/empty row that is not placed next to a sample row (green), exhibits a 10× reduced background luminescence. This spill-over of luminescence signal in in-between sample rows (light blue) translates to an 1% increase in remaining GTP - a small but unnecessary error to the assay’s accuracy.

### Supplement S6

**S6 Figure 1.**
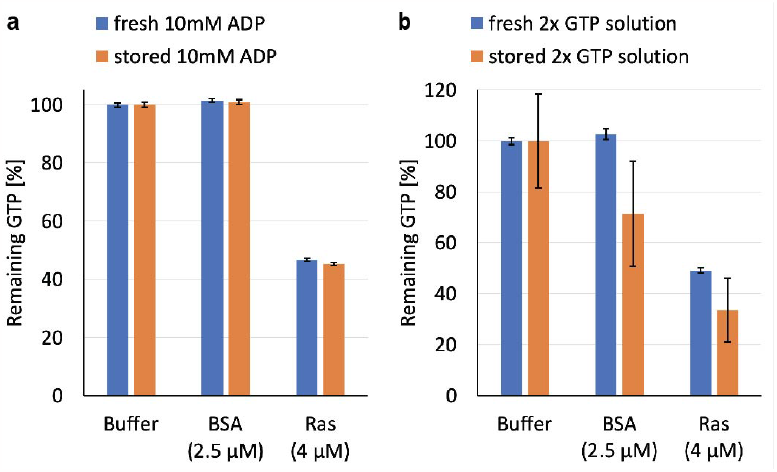
In GTPase assays the by the kit provided 10 mM ADP solution can be re-used (here: 3 freeze/thaw cycles) (a). In contrast, the 2× GTP solution can not be stored (b)! The graphs show the amount of remaining GTP for buffer (used for normalisation), BSA, and Ras GTPase. BSA is not a GTPase and does not change the GTP content. Ras is a GTPase and hydrolyses GTP, decreasing the amount of remaining GTP. Re-using 2× GTP solution results in huge variations between replicas of the same sample, leading to large error bars.

### Supplement S7

We advice to aliquote the detection reagent to reduce the number of freeze-thaw cycles and decrease the time required for thawing. Before using the detection reagent in GTPase assays, prepare a sufficient volume (e.g. through mixing of several aliquots) and vortex for proper mixing. We strongly advice against using separate detection reagent aliquots in one assay. *In some cases* this results in a large shift in luminescence, negating assay reliability (S7 Fig. 1).

**S7 Figure 1.**
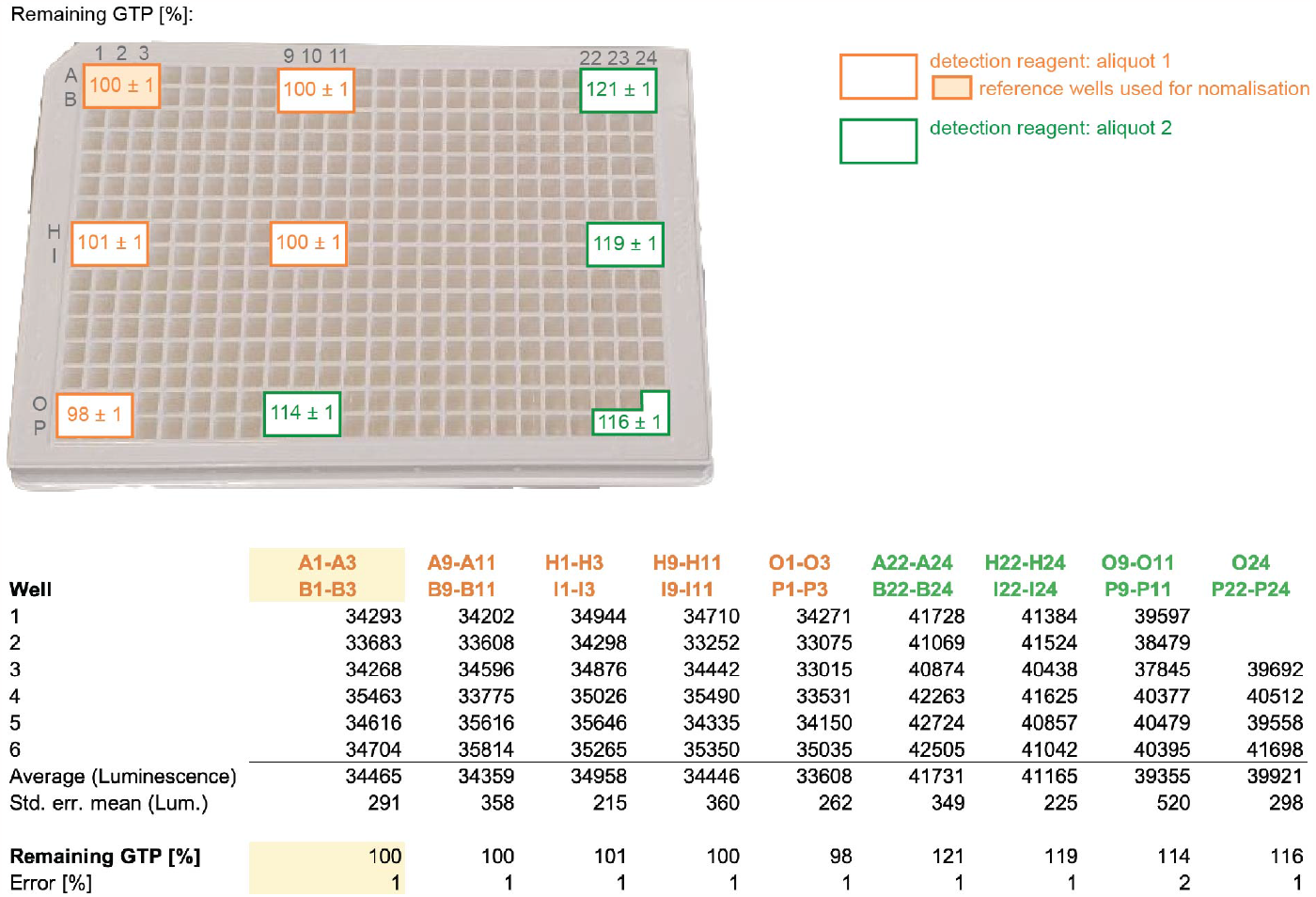
Luminescence (and resulting remaining GTP) values of a GTPase assay where buffer was added to all wells. In wells marked in orange one detection reagent aliquot was used and in wells marked in green a separate detection reagent aliquot was used. The use of distinct aliquots resulted in this case in a large difference in luminescence between both groups, propagating to an up to 20% difference in the perceived amount of remaining GTP.

### Supplement S8

We developed a GTPase activity model for determining the GTPase cycling rates *k*. It is briefly described in the following. (An extended version is given in S11.)

#### A GTPase cycling model

GTPase cycling involves three steps: (1) A GTP molecule from solution binds to the GTPase. (2) The GTPase hydrolyses GTP. (3) The GTPase releases GDP.

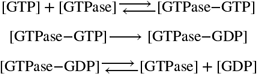

The activity of some GTPases can further be upregulated by effector proteins: GAPs have been shown to enhance GTP hydrolysis by the GTPase (step 2), GEFs enhance the release of GDP from the GTPase (step 3) [Bos et al., 2009, Vetter and Wittinghofer, 2001, Cherfils and Zeghouf, 2013].

To quantitatively describe the GTPase reaction cycle, we coarse-grained the GTPase reaction steps with

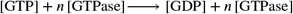

To account for possible GTPase dimerisation and cooperativity, we included the following reactions into the model:

1. Some GTPase enzymes can dimerise [Zhang and Zheng, 1998, Zhang et al., 1999, Zhang et al., 2001, Kang et al., 2010]:

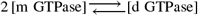

and both monomeric and dimeric forms of the GTPase can contribute to the overall GTP hydrolysis with different rates:

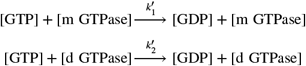

Assuming that the majority of the GTPase enzyme is in its monomeric form ([m GTPase] < *C*_*d*_, with *C*_*d*_ as the concentration at which half of the total GTPase is dimeric), we can approximate

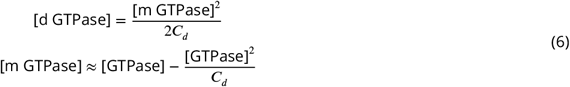
2. Next to cooperativity from dimerisation, cooperativity can also emerge when GTPase proteins come in close contact with each other - they can affect each other’s behaviour without forming a stable homodimer, effectively functioning as an effector protein for themselves:

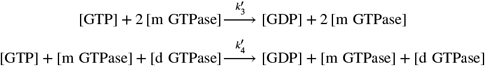
3. Effector proteins, such as GAPs and GEFs, affect the speed of the GTP hydrolysis cycle:

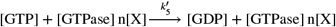

Here *X* is an effector protein and *n* ∈ ℕ.

Our data showed that the amount of remaining GTP follows an exponential decline over time (S4):

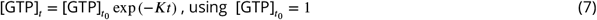

Considering reactions (1) - (3), we can thus define *K* in Eq. 7 as

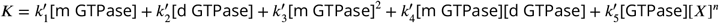

Using Eq. 6, and considering only up to second-order terms, results in

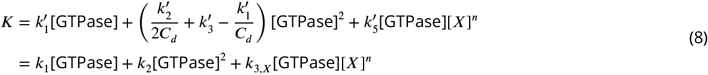

where *k*_1_ refers to GTP hydrolysis cycling rates of monomeric GTPase, *k*_2_ includes effects of cooperativity and dimerisation and *k*_3_ represents the rate of GTPase - effector interaction. We refer to *K* as ‘overall GTP hydrolysis rate’.

### Variability between assays

Eq. 8 with [X]=0 can be used to determine the rates of the GTPase alone. Then assays with the GTPase and an effector protein can be conducted to determine *k*_3_. While doing so one needs to account for assay variability, i.e. for the observation that the rates for the GTPase can vary between assays. Possible reasons for this include small concentration differences introduced though pipetting of small volumes (as are required for this assay), temperature and shaker speed fluctuations during the incubation step, and/or intrinsic changes in the protein activities due to other external conditions. To account for this variance, we introduced the parameter *c*_*corr*_. It maps all factors that lead to variations between assays onto the GTPase concentration.

The assay data, including samples containing only GTPase and GTPase - (effector *X*) mixtures, are fitted with

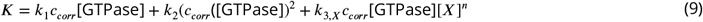

to determine *c*_*corr*_ and *k*_3,*X*_ (using *k*_1_ and *k*_2_ determined earlier) (with *n* either 1 or 2).

*c*_*corr*_ values are usually close to 1.0 (e.g. [Tschirpke et al., 2023b, Tschirpke et al., 2023a]), showing that the variation between assays is small. We advice to exclude assays with a big or very small *c*_*corr*_, as these indicate that the GTPase behaviour/assay conditions are unusual.

### GTPase - effector interactions

The accompanying matlab code allows to fit GTPase - effector mixtures that depend either linearly (*n* = 1) or quadratically (*n* = 2) on the effector concentration [*X*] (Eq. 9). If the effectors show neither a linear nor a quadratic concentration-dependence (e.g. due to saturation), we advice to either only include the linear/quadratic regimes into the analysis or extend our fitting model to match the specific case.

The model allows to fit GTPase - effector mixtures with up to two effectors present:

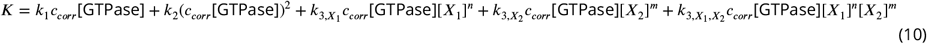

with *n* and *m* either 1 or 2.

#### Pooling of cycling rates *k* and error propagation

The way rate values are weighted for pooling and how errors are propagated is explained in detail in S11.

### Supplement S9

This supplement describes a simple python script that reads in GTPase data analysed in a spreadsheet editor (Support Protocol 1) and re-formats it into the input required for Support Protocol 2.

The script is illustrated using data of example 1 (Fig. 5).

#### Necessary resources

- Python script file: ‘Ras_example.ipynb’
- Data file: ‘example1.xlsx’
- a spreadsheet editor
- software to run a python script

#### Steps

- Open ‘Ras_example.ipynb’, state the input data and relevant tab names:

~~~
datafilename = ‘example1 . xlsx ‘
tabnamelist = [ ‘E1 ‘, ‘E2 ’ ]
~~~
- Run the python script. It will generate two excel sheeets: ‘E1.xlsx’ and ‘E2.xlsx’.
- Copy data of both outputs into one excel sheet, but only include one header (S9 Fig. 1)! This will be the input for the matlab script used in Support Protocol 2.
- Use the find/replace option of the spreadsheet editor to replace ‘.’ (a point) with ‘,’ (a comma).

*The python script generates numbers of the format ‘1*.*00’ while the matlab script requires the format ‘1,00’*.

#### General considerations on how the script operates

The spreadsheet data needs to conform to the following formatting to be processed by the python script (S9 Fig. 2):

- The script processes values in the spreadsheet area A80-Z89. *This area can only contain relevant numbers. If comments are placed in this area, the script will given an error*.
- The incubation time, stated in hours, must be stated in cell C82 (S9 Fig. 2 blue box).
- The error of the buffer must be stated in cell E81 (S9 Fig. 2 blue box).
- Remaining GTP values must be stated in cells F80-Z80. Cells not in use must remain empty (S9 Fig. 2 red box).
- Remaining GTP error values must be stated in cells F81-Z81. Cells not in use must remain empty (S9 Fig. 2 red box).
- Protein names must follow the formatting `’ProteinName_conc’` and be stated in A83-A89. Cells not in use must remain empty (S9 Fig. 2 orange box).
- Protein concentration values must be stated in the area B83-Z89. It is important to state the concentration of each protein listed in the protein name section here. (I.e. if a protein is not part of a sample, its concentration is 0.) Cells not in use must remain empty (Fig. 2 orange box).

**S9 Figure 1.**
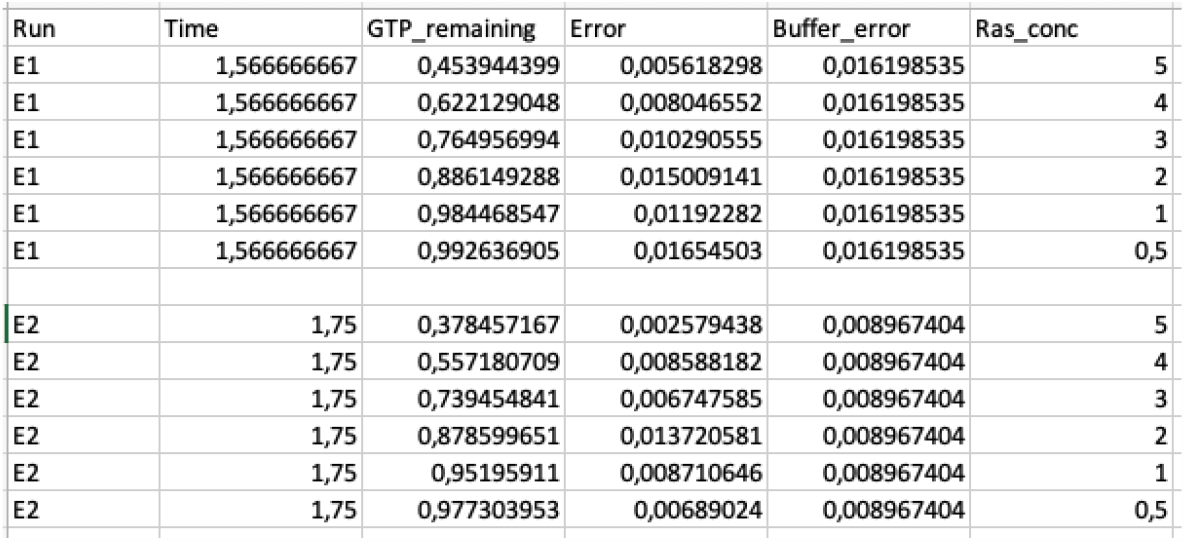
Required input format for the matlab scirpt (Support Protocol 2).

**S9 Figure 2.**
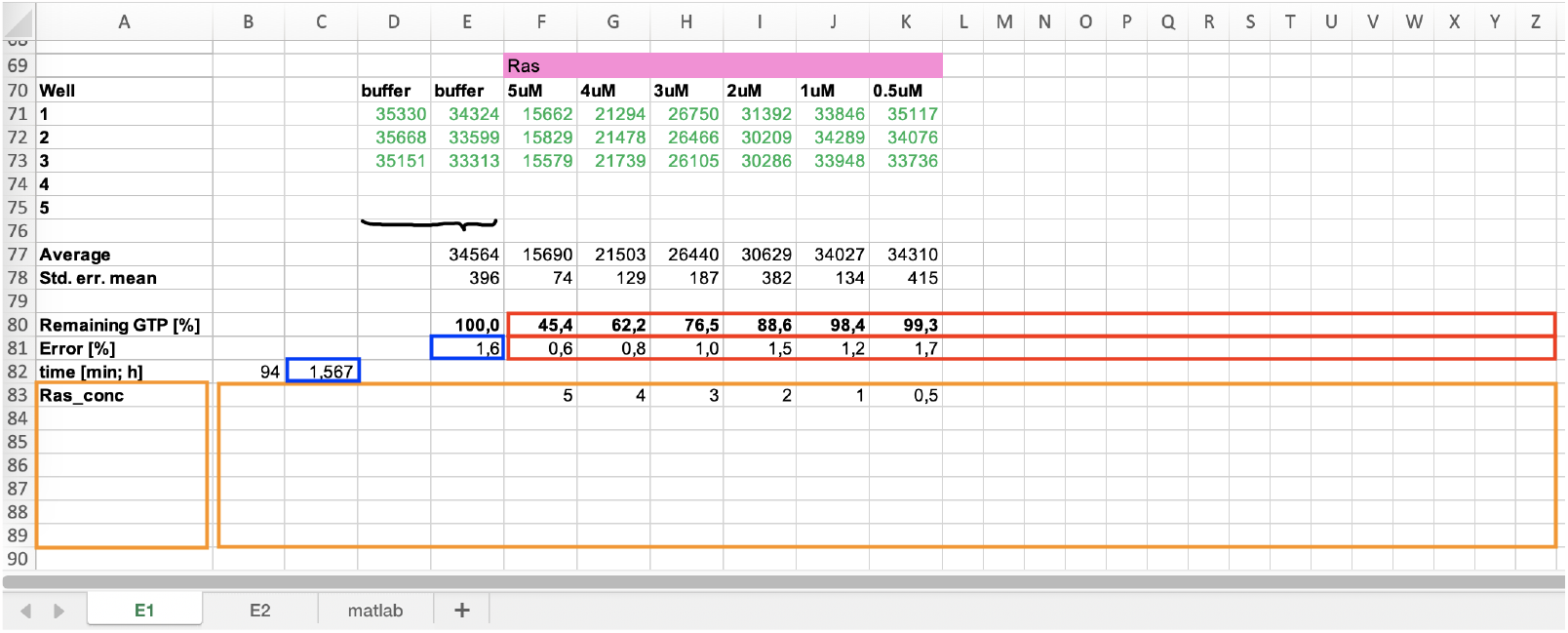
Required input format for the python scirpt.

### Supplement S10

To use the plotting scripts, a ‘Data_assays.mat’ file (output of Support Protocol 2) is required. All scripts automatically save the plot as ‘.tif’ and ‘.pdf’.

**S10 Figure 1.**
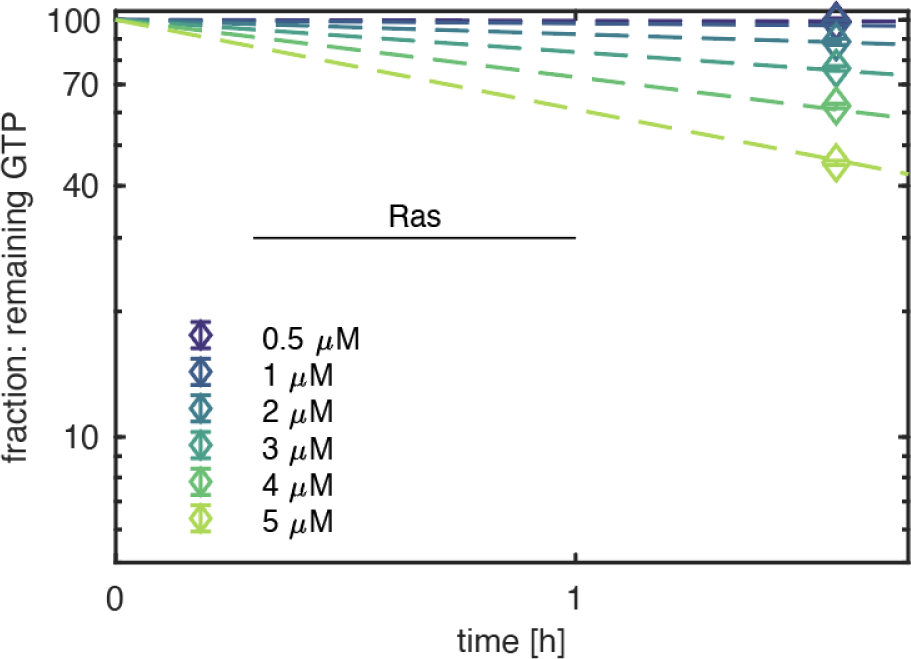
Exponential fit for the amount of amount of GTP over time for Ras concentrations. Generated for assay name ‘Ras’ and experiment number ‘E1’ (using ‘Plot_Semilog_GTP_time.m’ and ‘Data_assays.mat’ in the folder ‘example1and2 matlab output’).

#### Plot_Semilog_GTP_time.m

This script produces plots showing of the amount of GTP over time fitted with an exponential, as shown in S4. It can be used for assays of GTPases and GTPase effector mixtures with one effector.

1. Run ‘Plot_Semilog_GTP_time.m’.
2. Select a ‘Data_assays.mat’ file.
3. Choose an assay name, as stated in the previously used ‘assaylist.xlsx’ file. Then choose which assay (shown by assay number) should be plotted. An example plot is given in S10 Fig. 1. *This script is especially useful to assess/ verify that the amount of GTP declines exponentially. To do so, several assays (e*.*g. 3) of different incubation times using the same protein concentrations need to be conducted. To plot all (3) assays in the same plot, give these assays* ***the same*** *assay number before analysing them using ‘Process_assays*.*m’*.

**S10 Figure 2.**
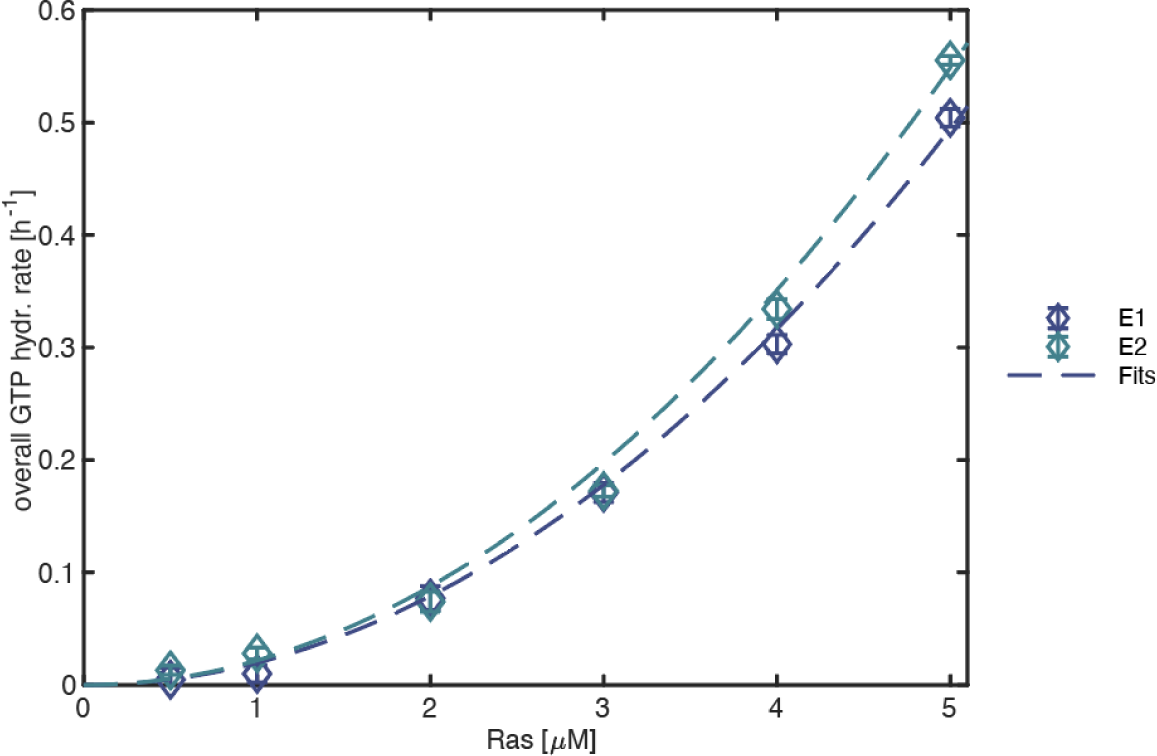
Overall GTP hydrolysis rate over Ras concentration for assays ‘E1’ and ‘E2’. Generated for assay name ‘Ras’ (using ‘Plot_rate_concentration.m’ and ‘Data_assays.mat’ in the folder ‘example1and2 matlab output’).

#### Plot_rate_concentration.m

This script produces allows to plot of the overall GTP hydrolysis rate for several assays in one figure. It can be used for assays of GTPases and GTPase effector mixtures with one effector.

1. Run ‘Plot_rate_concentration.m’.
2. Select a ‘Data_assays.mat’ file.
3. Choose an assay name, as stated in the previously used ‘assaylist.xlsx’ file. An example plot is given in S10 Fig. 2.

**S10 Figure 3.**
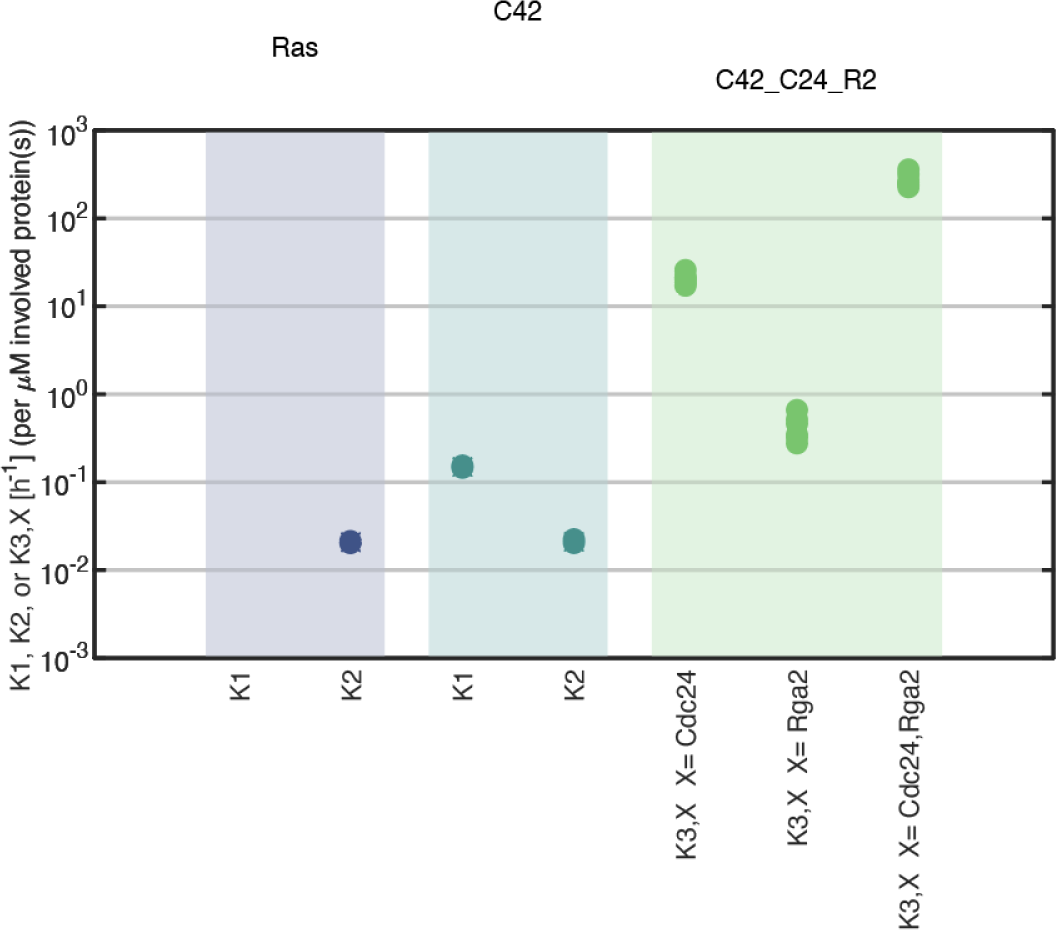
Plot of rates *K*_1_, *K*_2_ for Ras (left), *K*_1_, *K*_2_ for Cdc42 (middle), and *K*_3_, *X* for Cdc42-Cdc24-Rga2 assays. The assay name is stated on top. Rate values of individual experiments are shown as filled dots. The average is shown as a cross and error bars represent the standard error. Generated for assay names ‘Ras’, ‘C42’, and ‘C42-C24-R2’ (using ‘Plot_pooled_values_plot_std_err.m’ and ‘Data_assays.mat’ in the folder ‘example1and2 matlab output’).

#### Plot_pooled_values_plot_std_err.m

This script produces plots of the pooled rates *k*_1_, *k*_2_ *k*_3_ (that are also shown in the ‘Data_summary.xlsx’ file).

1. Run ‘Plot_pooled_values_plot_std_err.m’. *The parameter ‘y_limits’ in the code can be used to modify the limits of the y-axis. It is currently set to [1e-3 1e3]*.
2. Select a ‘Data_assays.mat’ file.
3. Choose an assay name, as stated in the previously used ‘assaylist.xlsx’ file. An example plot is given in S10 Fig. 3.

**S10 Figure 4.**
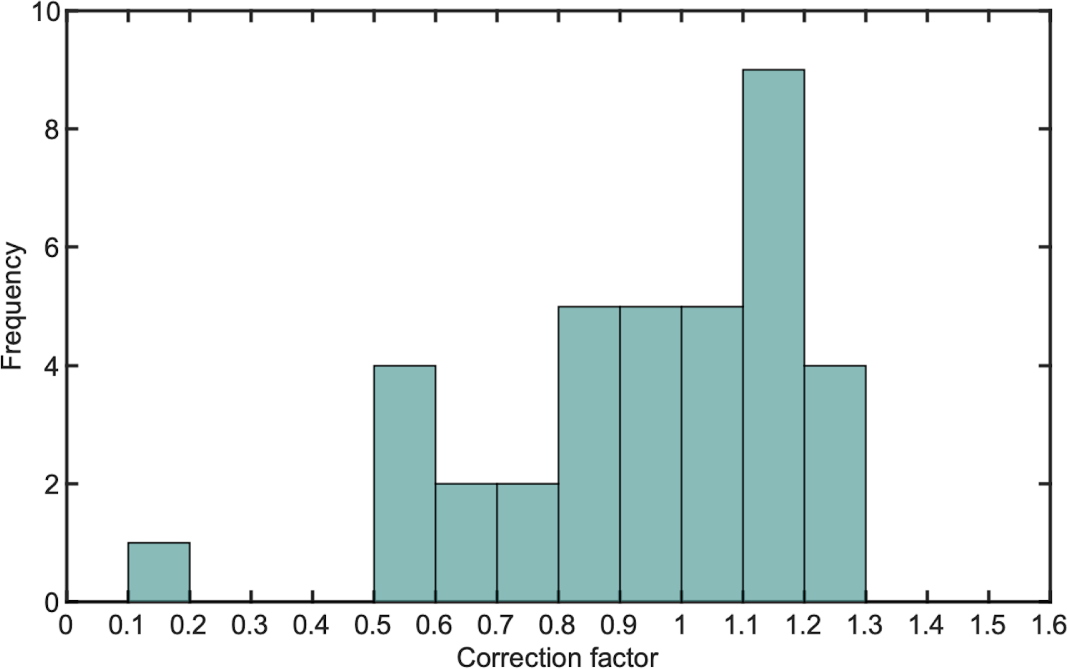
Histogram plot of *c*_*corr*_ using a bin size of 0.1. Generated for all *c*_*corr*_ values of example 2 (using ‘Plot_c_corr_histogram.m’ and ‘example2-histogram.xlsx’ in the folder ‘example2 matlab output’).

#### Plot_c_corr_histogram.m

This script produces a histogram plot of *c*_*corr*_ values. Before running the script, first copy/paste all *c*_*corr*_ values that should be plotted into a spreadsheet file.

1. Copy/paste all *c*_*corr*_ values that should be plotted from ‘Data_summary.xlsx’ into the first column of a spreadsheet file. All values should be in column A. Change the number formatting from ‘,’ to ‘.’ (i.e. change the number formatting from ‘0,863’ to ‘0.863’.
2. Run ‘Plot_c_corr_histogram.m’. An example plot is given in S10 Fig. 4. *The parameter ‘bin_size’, ‘x_start’, and ‘x_end’ in the code can be used to modify the plot. ‘bin_size’ states the bin size for the histogram segmentation, and ‘x_start’ and ‘x_end’ define the x-axis limits (i*.*e. minimum and maximum c*_*corr*_ *values that will be plotted)*.

#### Supplement S11

Crocodile stands for Crowding, cooperativity and dimerization in luminescence experiments. The purpose of this model is to describe, dissect and interpret the results of GTPase assays with, also in combination with other effectors, for example effector Cdc24 in the case Cdc42 is the GTPase. In these assays, GTP is hydrolyzed over time by a GTPase at a rate dependent on the concentration of proteins involved. The following section describe how we model the rate and what assumptions underlie this description.

## General model outline

We consider a GTPase in solution with nucleotides that get hydrolyzed through GTPase cycles. Effectors may also be present to speed up (parts of) the GTPase cycle. Chemically, the GTPase cycle consists of three steps, which are nucleotide binding, hydrolysis and nucleotide release, and effectors influence the rates of one or more of these steps. First considering monomeric GTPases with concentration [*m GTPase*], possible in complex with a nucleotide [*m GTPase* − *GNP*], this cycle constitutes the following reaction schemes (with GNP representing a GTP or GDP nucleotide):

Nucleotide binding (reaction rate constant 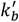 )

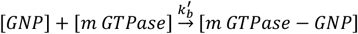

Hydrolysis (reaction rate constant 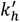] )

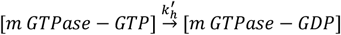

Nucleotide release (reaction rate constant 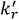 )

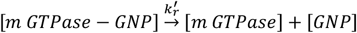

Effectors will influence the reaction rates and to avoid making concrete assumption on the molecular mechanism of each effector, and to reduce the number of fitting parameters later, we coarse-grain this GTPase cycle to a single step with rate constant 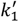. This will also help us to deal with the rate variability across replicate experiments as we will see further on.

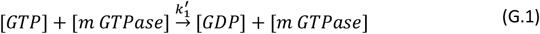

GTPases may also dimerize. To take this into account, we also consider the possible reaction (with rate constant 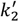 ):

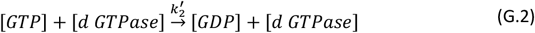

where GTPase dimers results from the monomers through the reaction:

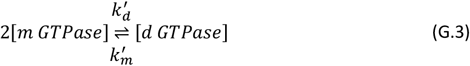

with monomeric and dimeric rate constants 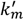 and 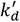 respectively.

When involving another protein X into this coarse-grained cycle such as an effector, we have (with rate constant 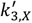 :):

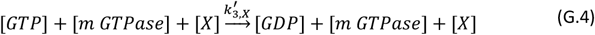

Generally, the overall uncorrected hydrolysis rate *K*^∗^ will then take the form (correction explanation follows later):

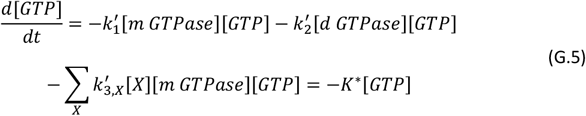

adding all contributions of each individual cycle reaction to the overall hydrolysis, potentially having multiple proteins *X* that contribute to the summation. This equation retains the same form if we instead assume complex formation between *X* and the GTPase, such that we have 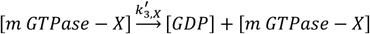: . This alternative leads to:

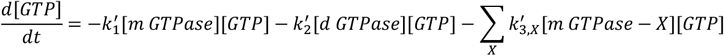

but as we expect [*m GTPase* − *X*] ∝ [*m GTPase*][*X*] from the rate equation of [*m GTPase*] + [*X*] ⇄ [*m GTPase* − *X*] in equilibrium, this is still equivalent to G.5.

Assuming the GTP concentration contains the only time-dependence on the right-hand side (i.e., all proteins have had time to equilibrate their reaction with each other), we can write:

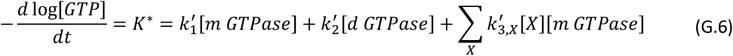

and this yields an exponential decay for the GTP nucleotides:

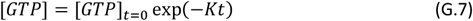

### Pooling

Generally, we have multiple runs for an experiment with a GTPase, each yielding estimates for e.g., *k*_1_ and *k*_2_ with standard errors. To get a single estimate, we must pool these. This is done by performing a weighted average to obtain the pooled estimates, as explained in the Supplements of (Tschirpke, Daalman & Laan, 2023).

In short, we model for example the individual run estimates 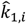 (with *i* = 1, 2, …, *n* with *n* as the number of runs) to deviate from the pooled *k*_1,*p*_ through normally distributed errors, but with heteroscedasticity:

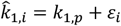

with errors *ε*_1_ ∼ 𝒩(0, *σ*_1_) and *σ*_1_ as the standard errors of each run estimate. By multiplying both sides by the weights *w*_*i*_ = 1*/σ*_*i*_, the resulting weighted estimates per run now follow a standard normal distribution. As a consequence, estimates that have large uncertainty are awarded a lower weight for the weighted estimated of *k*_1,*p*_ that follows. This weighted estimate of *k*_1,*p*_ is constructed by minimizing the (sum of) weighted errors, realized by squaring these first to treat positive and negative errors equally. This minimization amounts to a weighted least squares regression to yield the estimate 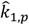 of *k*_1,*p*_, see e.g., (Heij, de Boer, Franses, Kloek, & van Dijk, 2004), which is a weighted average of the run estimates:

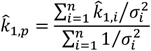

with standard error:

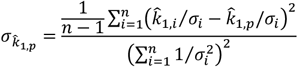

By the same token, we can construct other pooled estimates such as 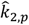.

## Case 1: No effectors

Even in absence of effectors, reactions G.1 and G.2 is not necessarily the only reaction taking place. GTPase can exhibit cooperativity and/or dimerization, Thus, we also consider reaction G.4 with *X* as another GTPase molecule, and this reaction reduces to (with rate constant 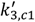 ):

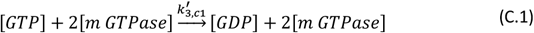

which implies cooperativity. Presence of dimers can also be taken into account with X = d GTPase, so we also have (with rate constant 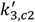 ):

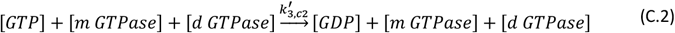

Theoretically, even higher order encounters (e.g., dimer + dimer) of GTPase molecules may take place, but we assume these get progressively unlikelier.

Given the previous reactions, the overall uncorrected cycling rate G.6 for the change in GTP concentration over time becomes (with *X* = {[*m GTPase*], [*d GTPase*]}:

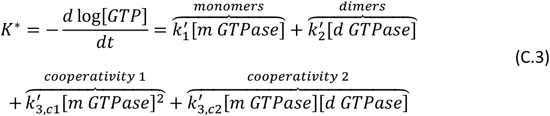

The overall cycling rate is hence a function of the monomeric and dimeric GTPase concentration. However, we supply a specific total GTPase concentration, so we want to rewrite all terms to this known [*GTPase*] concentration. Assuming the monomeric and dimeric pool are in equilibrium with each other, the rate equation resulting from the reaction in G.3 reads

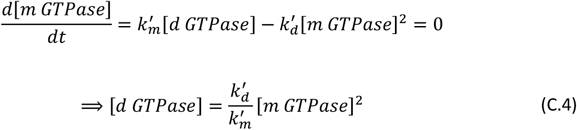

We can thus write the Hill equation:

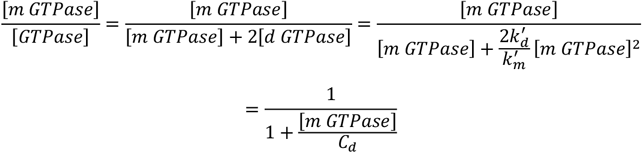

Consequently, C.4 then reads as:

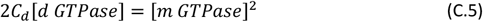

with 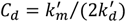. So, there is a critical concentration above which the monomeric fraction is low compared to the dimeric fraction. This critical concentration is higher if monomerization reaction rate 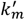 is high relative to dimerization reaction rate 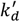.

Therefore, we can write the monomeric concentration as:

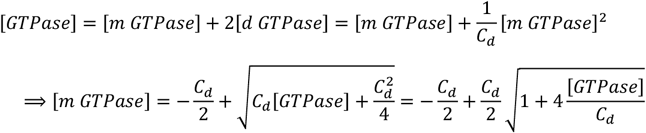

We can then use Taylor expansion to approximate this square root. The order up to which we need to expand depends on the GTPase concentration relative to the critical concentration, i.e. the size of the monomeric fraction relative to the dimeric fraction. If [*GTPase*] ≪ *C*_*d*_, then most GTPase molecules are monomeric and expansion up to second order (around the point 4[*GTPase*]/*C*_*d*_ = 0) suffices. As our data shows these terms suffices for fitting the GTPase dilution data well, we continue the derivation with the second order expansion. If other GTPase dilution data shows signs of higher order [*GTPase*] dependencies, this derivation must be adapted to include higher order expansion terms of the square root.

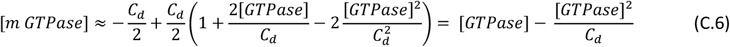

Similarly, using [*GTPase*] = [*m GTPase*] + 2[*d GTPase*] and substituting E.4, we get

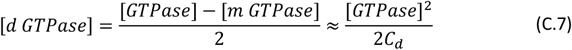

and also, combining E.3 and E.5, we get:

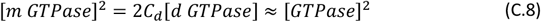

This means for the hydrolysis rate C.3, using the previous three equations:

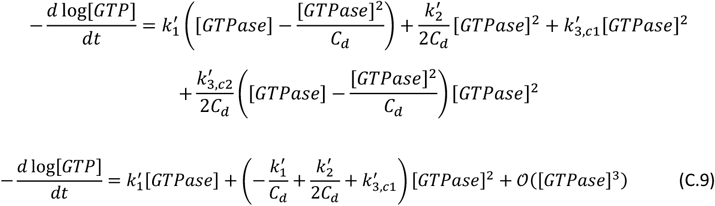

Ignoring the higher order terms 𝒪([*GTPase*]^3^) and defining 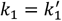 and 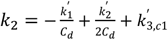, we obtain:

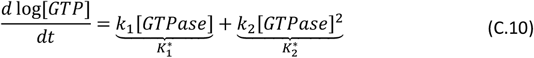

where *k*_1_ and *k*_2_ are cycling rates and 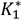 and 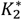 are uncorrected overall rate contributions to *K*^∗^. The former involves monomeric GTPase contributions, and the latter also <u>co</u>operativity and <u>di</u>meric contributions. In the section on optionally adding crowding effects, we see that <u>cro</u>wding would also present itself in the second term, explaining the origin for the ‘crocodile’ model term.

So, using G.7 we obtain for the GTP concentration:

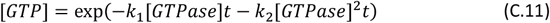

## Case 2: Adding a single effector

When an effector is added to the GTPase, more terms result from G.6 than those in C.11. Assuming the number of monomers and dimers is not significantly affected by the presence of GTPase-effector complexes, the same derivation as without effectors applies from G.6 up to C.9, only with an added term:

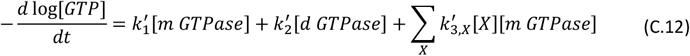

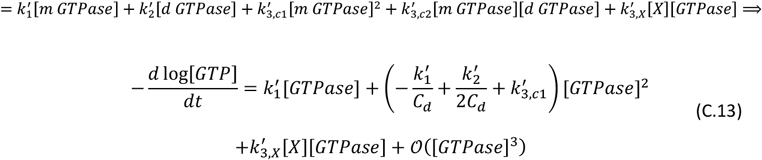

Defining 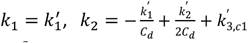 and 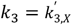, and ignoring the higher order terms 𝒪([*GTPase*]^3^) leads to:

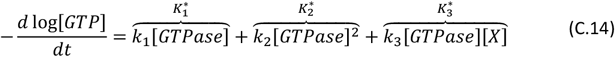

The new cycling rate *k*_3_ and uncorrected overall rate contribution 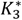 reflect the interaction between effector and GTPase. Note that a low value of *k*_3_ may be interpreted as absence of interaction, but can also be due to low functionality of the effector, or that the effector accelerates a step in the GTPase cycle which was already fast relative to the others. As we coarse-grain the full GTPase cycle into one step, the overall effectivity of the protein them appears low.

### GTPase concentration correction factors

In principle, *k*_1_ and *k*_2_ are known from an GTPase serial dilution assay (without the effector). However, in practice a complication arises. Expected activity of the GTPase may vary slightly across experiments, for many reasons. This can be due to concentration variability introduced by pipetting, environmental variability from the ideal protocol situation or protein integrity variability. Moreover, the amount of GTPases molecules sequestered by complexes with the effectors may be non-negligible as previously assumed, which would lead to a lower effective monomer and dimer concentration and thus to a lower contribution of terms involving *k*_1_ and *k*_2_ than expected from the assay without effectors.

A modelling solution is to introduce a run-specific GTPase concentration correction factor for assays with effectors, to account for these run-specific effects. This factor is a single constant applying to all instances of [*GTPase*]. The correction factor also provides us with a diagnostic for run-specific issues, as ideally the value is ideally close to 1. Large deviations from 1 are indicative of incidental (e.g., a pipetting error) or systematic (e.g., strong sequestration of GTPases in complexes) problems in the assay (modelling). The code implementation of the model therefore allows the user to define what range of correction factors is still deemed acceptable, and those runs with correction factors that do not fall in this range are excluded for pooling single-run parameter estimates to the pooled estimates.

Including this correction factor *c*_*corr*_ into C.14, the corrected overall rate contribution *K* then becomes:

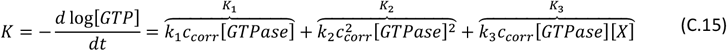

with *K*_1_, *K*_2_ and *K*_3_ as overall rate contributions. To fit this function, we still see it is of the form *a* + *b*[*X*], where *k*_3_ is contained in *b*. But before we can retrieve *k*_3_, we must determine *c*_*corr*_. This can be done by comparing the overall cycling rate *K* of the effector assay when [*X*] = 0, to the GTPase dilution assay cycling rate (which has no effector). Concretely, we compare:

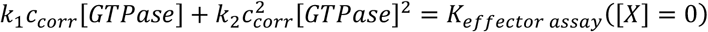

The [*GTPase*] is set to the typical value in the effector assay. The left-hand side uses the *k*_1_ and *k*_2_ estimates that have been previously established from the GTPase serial dilution assay, while the right-hand side is the at [*X*] = 0.

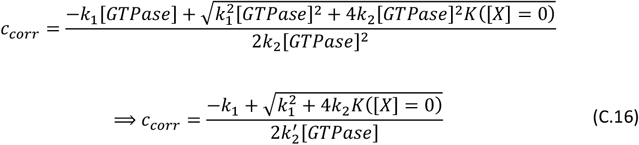

The rate of the effector assay can be measured directly or more reliably, inferred from evaluation at [*X*] = 0 of the fit based on all effector assay points, as in the computational implementation of the model. More specifically, this implementation generates random draws based on the fitting errors, while the fit also takes errors on the data points into account. Consequently, we can generate not only the point estimate of the correction factor, but also with the standard error. However, the code only uses the point estimate to excessively avoid inflating the errors on subsequent rate parameter estimates. For example, if we fit in practice *K* = *a* + *b*[*X*], then 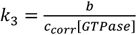.

### Linear and quadratic effectors

Some effectors form complexes with a GTPase, but can also dimerize themselves. If it can be assumed that this dimer is the most relevant form of the protein (due to its abundance or activity), this means that C.14 becomes (with *X* as dimer *Y*:*d Y*):

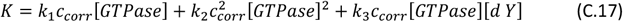

As dimer *Y* originates from two monomers, we have 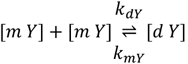, with the rate equation:

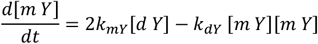

In equilibrium, this means [*d Y*] ∝ [*m Y*]^2^, and analogously to the GTPase monomer-dimer equilibrium, [*d Y*] ∝ [*Y*]^2^ (see (C.7)), such that *K* gets a quadratic dependence on effector concentration. The code implementation of the model accommodates both a linear dependency of the overall rate on effector concentration and a quadratic dependency. Which case applies per effector can be defined by the user.

## Case 3: Adding two effectors

With two effectors instead of one, the summation on the right hand side of C.12 will contain multiple terms for every effector combination. If we consider two effectors *X*_1_ and *X*_2_, the set whose rate contributions must be considered contains these two and a potential cross-term *X* = {*X*_1_, *X*_2_, *X*_3_ – *X*_2_}. As a result, C.12 will become:

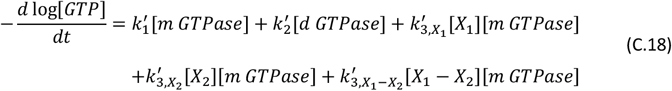

where we can replace [*X*_1_ – *X*_2_] with [*X*_1_][*X*_2_] as 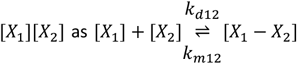 leads to rate equation 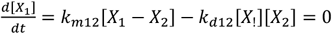 in equilibrium.

Proceeding analogously to the one effector case, ignoring the higher order terms 𝒪([*GTPase*]^3^) again, we obtain:

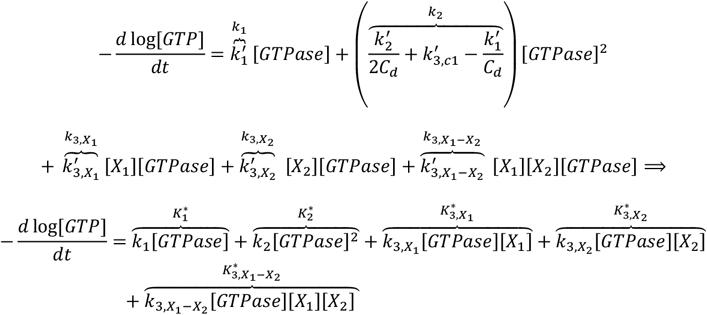

Taking into account the [*GTPase*] correction factors, this becomes:

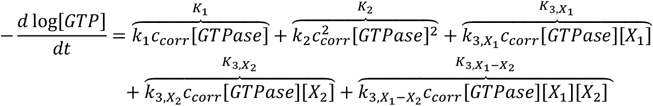

The last term represents an interaction term between effectors, indicating a possible synergy between the effects of the proteins on the GTPase cycle. As with the single effector case, the effectors can be linear or quadratic, such that e.g., for a quadratic effector *X*_1_ every instance of [*X*_1_] can be replaced by [*X*_1_]^2^ if *X*_1_ is known to dimerize into a more active form and/or to be abundant in dimers.

## Fitting restrictions on parameters

Summarizing the three cases of 0, 1 and 2 effectors, we have seen that the rate equations read respectively:

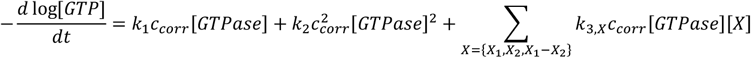

where *c*_*corr*_ = 1 for the zero-effector case. This means we have 2, 4 or 6 parameters in the 0, 1 and 2 effector cases respectively.

However, when fitting this function, we must also take into account restrictions on the parameters we know must exist. A GTPase by default must contribute positively to hydrolysis, So the main term with *k*_1_ must be positive, i.e. *k*_1_ > 0. As the ‘crocodile term’ *k*_2_ might include multiple effects such as crowding, we do not impose this restriction on *k*_2_. Similarly, for assay with effectors where only the effector concentration is varied, the equation takes the form of *a* + *b*[*X*_1_] + *c*[*X*_2_] + *d*[*X*_1_][*X*_2_], where we assume the contribution *a* of the GTPases without effector must be positive.

For this purpose, the code implementation of the model transforms all *k*’s to an alternative parameter space. Concretely, we define:

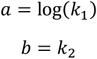

when fitting the no effector case:

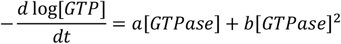

When fitting with effectors, we define:

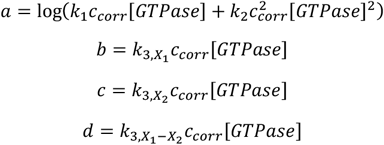

when fitting the effector case:

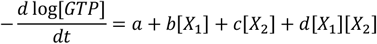

The individual rate constant estimate of *k*_3_’s can then be retrieved by dividing out the *c*_*corr*_ obtained as explained in the GTPase concentration correction factor section in Case 2: Adding a single effector.

Moreover, we restrict the concentration correction factor to the range 0.1 to 10 to avoid an apparent e.g., negative activity or extreme overactivity of a GTPase in a particular assay. Runs hitting these bounds for *c*_*corr*_ are indicative of a possible issue.

## Error propagation

Ultimately, we obtain pooled estimates of cycling rates *k*_1_ and *k*_1_ and if applicable, *k*_3_’s which also has standard errors as explained in the pooling section in the General model outline. Originally, we have errors on the measurements of the GTP concentration. The squared reciprocals of these errors provide the weights for the non-linear regression in Matlab’s fitnlm. We then generate random draws of the fitting parameter values in the alternative parameter space (see section Fitting restrictions on parameters) through multivariate normal random variables with the parameter covariance matrix of fitnlm. These draws are then transformed back and if needed divided by the concentration correction factor and the [*GTPase*] to obtain the draws of the original parameters. Uncertainty in concentration correction factors are not propagated as mentioned in the section on these factors to not excessively inflate the rate parameter estimate errors. Finally, we use the standard deviations of these draws for the standard errors to use in the pooling regression as explained in the pooling section.

## Optional crowding addition

Sometimes, crowding effects between proteins may be expected to play a role in the reaction rates. These effects can be accommodated in the Crocodile model. From literature, it is not evident whether we should expect a positive or negative contribution of crowding to the reaction rate (Kim and Yethiraj, 2009). Therefore, we approximate this as the same linear dependency of all reaction rate constants *k*’, related to the GTPase cycle reactions, on the total protein concentration. This dependency can hence be positive or negative. Concretely, the rate constants then change through:

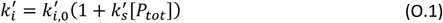

with [*P*_*tot*_] as the total concentration of proteins present and 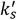 as a constant that can be positive, 0, or negative.

As we show below, we retain the same functional forms for the GTP hydrolysis rate. The only change is the way to interpret *K*_2_ and the *K*_3_’s (and concordantly, *k*_2_, the 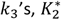 and the 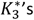), which now includes a crowding term. Incidentally, as *K*_2_ now encompasses <u>cro</u>wding, <u>co</u>operativity and <u>di</u>merization (of luminescence <u>e</u>xperiments), this can be dubbed the crocodile-term.

### Single effector case

We can insert crowding in C.13 through O.1 such that e.g., 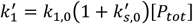 with [*P*_*tot*_] = [*GTPase*] + [*X*] and similarly for 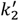 and 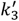. Ignoring as before the higher order terms 𝒪([*GTPase*]^3^), we get:

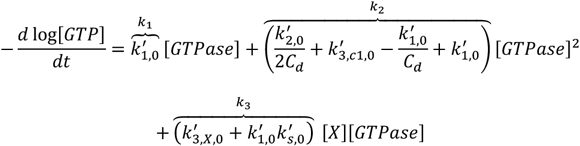

This is the same functional form as before in C.14, only the interpretation of *k*_2_ and *k*_3_ contains crowding.

### Double effector case

We follow the same logic starting from C.18, only this time incorporating crowding through O.1 with [*P*_*tot*_] = [*GTPase*] + [*X*_1_] + [*X*_2_]. Ignoring the higher order terms 𝒪([*GTPase*]^3^) yields:

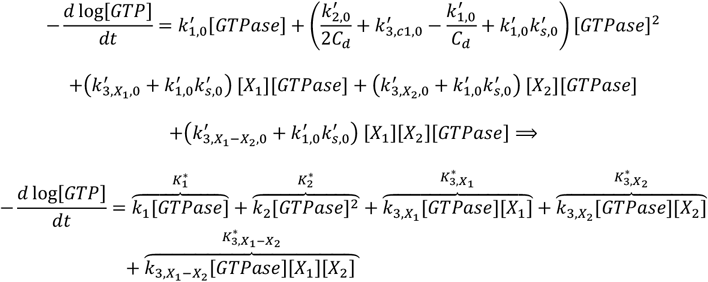

Once more, we obtain the same functional form as without crowding, only the interpretation of *k*_2_ and the *k*_3_’s is broadened.

## References

Bezeljak, U., Loya, H., Kaczmarek, B., Saunders, T. E., and Loose, M. (2020). Stochastic activation and bistability in a Rab GTPase regulatory network. Proceedings of the National Academy of Sciences, 117(12):6540–6549.

Bos, J., Rehmann, H., and Wittinghofer, A. (2009). GEFs and GAPs: Critical Elements in the Control of Small G Proteins. Cell, 16(3):374–383.

Cherfils, J. and Zeghouf, M. (2013). Regulation of small GTPases by GEFs, GAPs, and GDIs. Physiological Reviews, 93(1):269–309.

Kang, P. J., Béven, L., Hariharan, S., and Park, H. O. (2010). The Rsr1/Bud1 GTPase interacts with itself and the Cdc42 GTPase during bud-site selection and polarity establishment in budding yeast. Molecular Biology of the Cell, 21(17):3007–3016.

Kohyama, S., Merino-Salomón, A., and Schwille, P. (2022). In vitro assembly, positioning and contraction of a division ring in minimal cells. Nature Communications, 13(1).

Loose, M., Fischer-Friedrich, E., Ries, J., Kruse, K., and Schwille, P. (2008). Spatial Regulators for Bacterial Cell Division Self-Organize into Surface Waves in Vitro. Science, 320(5877):789–792.

Promega Corporation (2015). GTPase-Glo ™ Assay.

Rapali, P., Mitteau, R., Braun, C., Massoni-Laporte, A., Ünlü, C., Bataille, L., Arramon, F. S., Gygi, S. P., and McCusker, D. (2017). Scaffold-mediated gating of Cdc42 signalling ?ux. eLife, 6:1–18.

Ruxton, G. D. (2006). The unequal variance t-test is an underused alternative to Student’s t-test and the Mann-Whitney U test. Behavioral Ecology, 17(4):688–690.

Smith, N. J., van der Walt, S., and Firing, E. (2015). Magma, inferno, plasma and viridis colormaps.

Tschirpke, S., Daalman, W., and Laan, L. (2023a). The GEF Cdc24 and GAP Rga2 synergistically regulate Cdc42 GTPase cycling. BioRxiv.

Tschirpke, S., van Opstal, F., van der Valk, R., Daalman, W. K.-G., and Laan, L. (2023b). A guide to the in vitro reconstitution of Cdc42 activity and its regulation. BioRxiv.

Vendel, K. J. A., Tschirpke, S., Shamsi, F., Dogterom, M., and Laan, L. (2019). Minimal in vitro systems shed light on cell polarity. Journal of Cell Science, 132(4):1–21.

Vetter, I. R. and Wittinghofer, A. (2001). The guanine nucleotide-binding switch in three dimensions. Science, 294(5545):1299–1304.

Zhang, B., Gao, Y., Moon, S. Y., Zhang, Y., and Zheng, Y. (2001). Oligomerization of Rac1 GTPase Mediated by the Carboxyl-terminal Polybasic Domain. Journal of Biological Chemistry, 276(12):8958–8967.

Zhang, B., Wang, Z. X., and Zheng, Y. (1997). Characterization of the interactions between the small GT-Pase Cdc42 and its GTPase-activating proteins and putative effectors: Comparison of kinetic properties of Cdc42 binding to the Cdc42-interactive domains. Journal of Biological Chemistry, 272(35):21999–22007.

Zhang, B., Zhang, Y., Collins, C. C., Johnson, D. I., and Zheng, Y. (1999). A built-in arginine finger triggers the self-stimulatory GTPaseactivating activity of Rho family GTPases. Journal of Biological Chemistry, 274(5):2609–2612.

Zhang, B. and Zheng, Y. (1998). Negative regulation of Rho family GTPases Cdc42 and Rac2 by homodimer formation. Journal of Biological Chemistry, 273(40):25728–25733.

## Bibliography

Heij, C., de Boer, P., Franses, P. H., Kloek, T., & van Dijk, H. K. (2004). Econometric methods with applications in business and economics. Oxford University Press.

Kim, J.S., and Yethiraj, A. (2009). Effect of macromolecular crowding on reaction rates: a computational and theoretical study. Biophys. J. 96, 1333–1340.

Tschirpke, S., Daalman, W., and Laan, L. (2023). The GEF Cdc24 and GAP Rga2 synergistically regulate Cdc42 GTPase cycling. BioRxiv.

